# Evolution of ecological dominance of yeast species in high-sugar environments

**DOI:** 10.1101/020370

**Authors:** Kathryn M. Williams, Ping Liu, Justin C. Fay

**Author notes:** Corresponding Author: Kathryn Williams 4444 Forest Park Ave. St. Louis, MO 63108.

## Abstract

In budding yeasts, fermentation in the presence of oxygen evolved around the time of a whole genome duplication (WGD) and is thought to confer dominance in high-sugar environments because ethanol is toxic to many species. While there are many fermentative yeast species, only *Saccharomyces cerevisiae* consistently dominates wine fermentations. In this study, we use co-culture experiments and intrinsic growth rate assays to examine the relative fitness of non-WGD and WGD yeast species across environments to assess when *S. cerevisiae*’s ability to dominate high-sugar environments arose. We show that *S. cerevisiae* dominates nearly all other non-WGD and WGD species except for its sibling species *S. paradoxus* in both grape juice and a high-sugar rich medium. Of the species we tested, *S. cerevisiae* and *S. paradoxus* have evolved the highest ethanol tolerance and intrinsic growth rate in grape juice. However, the ability of *S. cerevisiae* and *S. paradoxus* to dominate certain species depends on the temperature and the type of high-sugar environment. Our results indicate that dominance of high-sugar environments evolved much more recently than the WGD, most likely just prior to or during the differentiation of *Saccharomyces* species, and that evolution of multiple traits contributes to *S. cerevisiae*’s ability to dominate wine fermentations.

## Introduction

Evolutionary innovation can promote the ecological dominance of some lineages by enabling them to occupy new niches. While conservation of an innovation among descendent taxa reflects its contribution to their ecological success, ecological dominance may not be an immediate consequence of evolutionary innovation. Phylogenetic studies indicate that the current dominance of some lineages may result from events temporally distinct from major evolutionary transitions (Wing and Boucher 1998; Alfaro et al. 2009; Edwards et al. 2010; Near et al. 2012; Schranz et al. 2012; Bouchenak-Khelladi et al. 2014). This apparent lag between the evolution of an innovation and the rise to dominance of descendent lineages may occur because dominance depends upon certain environments, ecological communities or the acquisition of additional traits.

In budding yeasts, evolution of the ability to ferment sugar in the presence of oxygen dramatically changed the way some species harness energy. Whereas most species acquire energy through respiration in the presence of oxygen, certain species such as *Saccharomyces cerevisiae* acquire most of their energy via the less efficient process of fermentation (Pronk et al. 1996). Evolution of this fermentative lifestyle likely involved multiple steps both before and after a whole genome duplication (WGD) in the yeast lineage, including the ability to grow without mitochondrial electron transport and the transcriptional rewiring of carbon metabolizing enzymes (Ihmels et al. 2005; Merico et al. 2007; Field et al. 2009; Hagman et al. 2013; Lin et al. 2013). While the evolutionary transition to a fermentative lifestyle began prior to the WGD, lineages that diverged after the WGD show a clear preference for fermentation in the presence of oxygen (Merico et al. 2007; Hagman et al. 2013).

Fermentation in the presence of oxygen is thought to provide WGD yeast species with a fitness advantage in high-sugar environments such as grape juice (Wolfe and Shields 1997; Piskur and Langkjaer 2004; Thomson et al. 2005; Piskur et al. 2006; Conant and Wolfe 2007). Theoretical modeling shows that a fermentative lifestyle can yield a growth advantage in high-sugar environments due to a higher rate of sugar consumption and energy production (Pfeiffer et al. 2001; MacLean and Gudelj 2006; Conant and Wolfe 2007). Additionally, ethanol produced during fermentation may inhibit the growth of competitor species (Gause 1934; Piskur and Langkjaer 2004; Thomson et al. 2005; Piskur et al. 2006). Thus, the fermentative lifestyle is expected to enable WGD species to dominate high-sugar environments like grape juice.

While *S. cerevisiae* has been shown to dominate competitions with multiple non-WGD species (Holm Hansen et al. 2001; Pérez-Nevado et al. 2006), the importance of the fermentative lifestyle remains equivocal. Competition experiments between *S. cerevisiae* and several non-WGD species did not support the role of ethanol but instead implicate different factors depending upon which competitor species was used.

Competitions with *Torulaspora delbrueckii* and *Lachancea thermotolerans* demonstrated that low-oxygen and cell-density contribute to *S. cerevisiae*’s dominance (Holm Hansen et al. 2001; Nissen et al. 2003, 2004), while competitions with *Hanseniaspora guilliermondii* and *H. uvarum* showed that *S. cerevisiae* produces a toxic metabolite derived from glyceraldehyde 3-phosphate dehydrogenase peptides (Pérez-Nevado et al. 2006; Albergaria et al. 2010; Branco et al. 2014). While *S. cerevisiae* exhibits high-ethanol tolerance (Pina et al. 2004; Belloch et al. 2008; Arroyo-López et al. 2010; Salvadó et al. 2011a), mono-culture growth rates of various species indicate that temperature is more important to *S. cerevisiae*’s dominance than ethanol tolerance (Goddard 2008; Salvadó et al. 2011a).

Within the vineyard environment, grapes and wine musts contain hundreds of yeast species, including a number of fermentative species (Pretorius 2000; Fleet 2003, 2008; Jolly et al. 2006; Bokulich et al. 2012; Pinto et al. 2014). Yet, even without the introduction of commercial wine yeast, *S. cerevisiae* consistently dominates grape juice as it ferments to wine (Fleet 2003, 2008). Since little is known about the relative fitness of most WGD species in high-sugar environments like grape juice, it is unclear whether *S. cerevisiae*’s dominance in wine fermentations reflects certain attributes of the grape juice environment or the yeast species present within the community, and whether dominance in high-sugar environments is a simple consequence of the fermentative lifestyle or involves the acquisition of additional traits.

The objectives of this study were to determine when the ability to dominate high-sugar environments evolved in the yeast lineage and to identify traits that confer *S. cerevisiae* with a growth advantage in these environments. To infer when dominance arose and lessen the impact of any potential strain or species outliers, we examined a taxonomically diverse sample of 18 different yeast species spanning the WGD and the evolution of the fermentative lifestyle (Figure 1). Given that the evolution of the fermentative lifestyle spanned the WGD (Hagman et al. 2013), we included non-WGD species that produce small amounts of ethanol (*Lachancea* and *Torulaspora* species) and WGD species that exhibit intermediate levels of fermentation (*Vanderwaltozyma* and *Tetrapisispora* species). To identify when dominance arose we directly competed these species with one another and found that dominance of high-sugar environments evolved much more recently than the evolution of the fermentative lifestyle. To identify traits involved in dominance we compared species’ intrinsic growth rates under a variety of conditions and found that both ethanol and the type of high-sugar environment influence fitness.

**Figure. 1.**
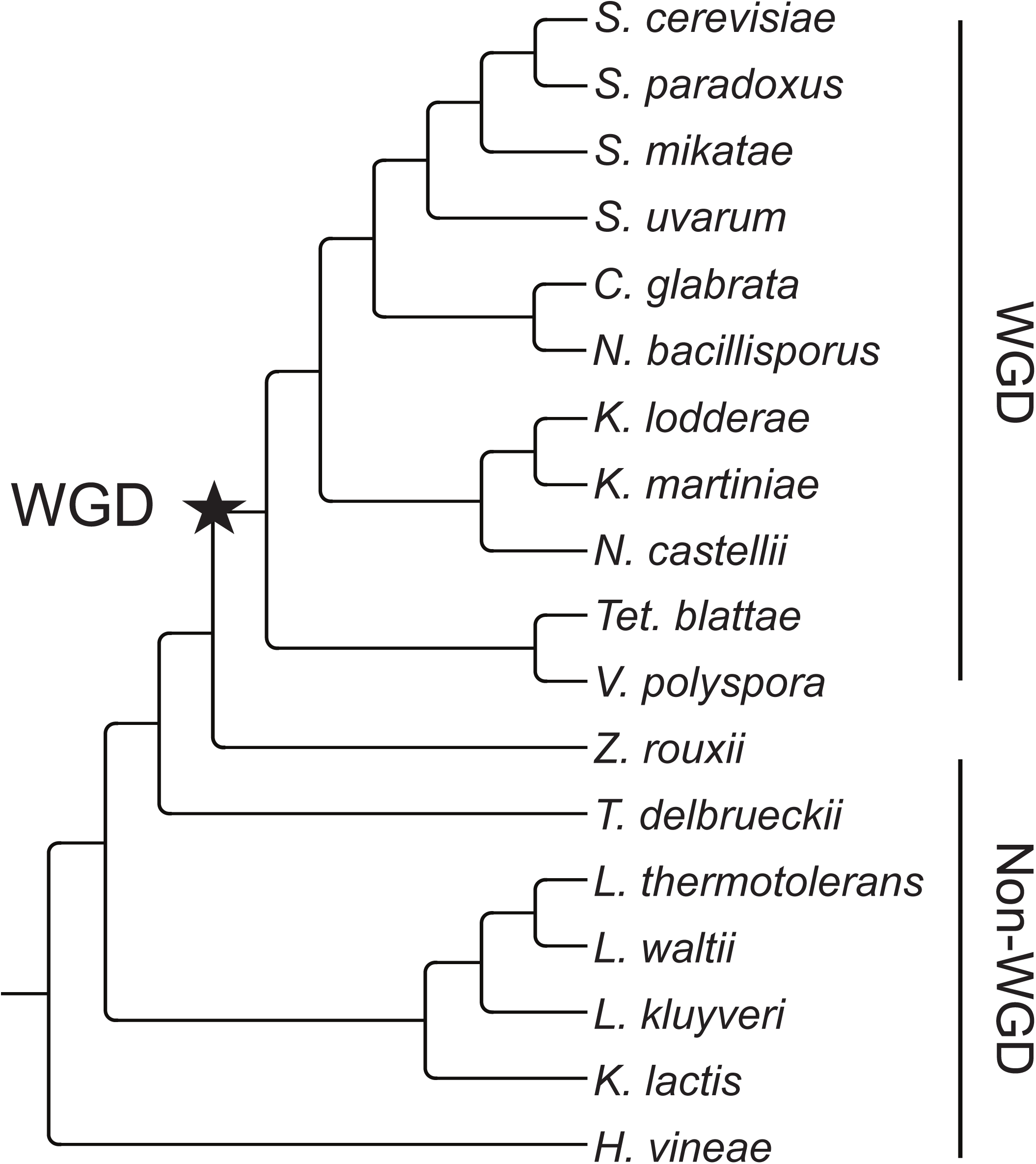
Phylogenetic relationship of yeast species used in this study. The phylogeny is based on two previous studies (Kurtzman and Robnett 2003; Salichos and Rokas 2013) and the placement of the *Nakaseomyces* (*C. glabrata* and *N. bacillisporus*) using chromosome rearrangements (Scannell et al. 2006). The whole genome duplication (WGD) event is shown by a star.

## Material and Methods

### Yeast strains

A total of 20 yeast strains representing 18 non-WGD and WGD species were used for our experiments (Table S1 and Figure 1). We chose a *S. cerevisiae* strain isolated from oak (YPS163) to represent *S. cerevisiae* (Sniegowski et al. 2002). As a control for population variation within *S. cerevisiae*, we also included I14, a *S. cerevisiae* strain isolated from a vineyard in Italy (Fay and Benavides 2005) and BJ20, a *S. cerevisiae* strain isolated from a tree in China (Wang et al. 2012).

### Growth media

The primary assay media used were two high-sugar environments: Chardonnay grape juice (Vintners Reserve, Winexpert Inc., Port Coquitlam, B.C., Canada), hereafter referred to as “Grape”, and high-sugar rich medium (10 g/l yeast extract, 20 g/l peptone, 120 g/l dextrose), hereafter referred to as “HS”. We chose HS to reflect the glucose concentration typical of the grape juice environment, ∼ 120 g/l (Rodicio and Heinisch 2009). Low-pH HS was made by adjusting the pH of HS from 6.7 to 3.7, the pH of our Grape medium, using tartaric acid, the predominant acid present in grape juice (Radler 1993). The media used to test for nutrient limitations in Grape included Grape with one of five nutrient supplements: YP (10 g/l yeast extract, 20 g/l peptone), CM (1.3 g/l synthetic complete with amino acids, 1.7 g/l yeast nitrogen base, and 5 g/l ammonium sulfate), AA (1.3 g/l complete amino acids), NB (1.7 g/l yeast nitrogen base), or AS (5 g/l ammonium sulfate). Assay media to test the ethanol tolerance of each yeast species was YPD (10 g/l yeast extract, 20 g/l peptone, 20 g/l dextrose) with ethanol concentrations ranging from 0-10%. Assay media to identify unknown inhibitor compounds produced by *S. cerevisiae* during growth was YPD made using supernatant from 16 other species (Table S1) grown in mono-culture and co-culture with *S. cerevisiae*. We chose YPD with 2% dextrose for ethanol tolerance and supernatant assays because ethanol produced during growth by fermenting species should not attain inhibitory concentrations.

### Competition experiments

#### Growth conditions

To assess the relative growth of non-WGD and WGD yeast species in high-sugar environments, we performed two competition experiments in which we measured the abundance of representative strains of multiple species relative to a focal species strain of *S. cerevisiae* or *S. paradoxus* after growth in co-culture. In the first competition experiment (Competition 1), we measured the abundances of six non-WGD and seven WGD species (Table S1) relative to a single *S. cerevisiae* strain (YPS163). As controls, we assessed the ability of the conspecific strain, I14, to grow relative to our reference *S. cerevisiae* strain, and grew each species in mono-culture. Competitions were initiated at approximately 10^3^ cells of each species per ml of Grape and HS media. One ml cultures were grown in 2 ml 96-well plates that were covered with breathable film and incubated at 30°C with shaking at 400 rpm for 48 hours. To assess the effects of strain and temperature on the relative growth of WGD species in high-sugar environments, we measured the abundance of five WGD species relative to three *S. cerevisiae* strains (YPS163, I14, and BJ20) and a single strain of *S. paradoxus* (YPS152) following co-culture (Competition 2). Cultures were prepared and grown as in Competition 1, except that replicates of each culture were grown at 30°C and 22°C.

#### Sampling

For both competition experiments, samples were taken at the beginning and the end of the experiment and frozen at -20°C for later use. For mono-cultures in Competition 1, cells from *S. cerevisiae* mono-culture were mixed in equal volume with cells from each of the other species’ mono-cultures at 48 hours and then frozen.

#### DNA extraction

DNA for Competition 1 was extracted from each sample using a protocol modified from Hoffman (2002) that included adding approximately 200 μl of 0.5 mm-diameter glass beads (BioSpec Products, Bartlesville, OK) to each sample and lysing cells in a bead beater (BioSpec Products) on high for 5 minutes at room temperature. For samples grown in Grape, DNA was also column purified to remove an unknown inhibitor of PCR amplification. DNA for Competition 2 was extracted using the ZR-96 Fungal/Bacterial DNA Kit (Zymo Research, Organge, CA).

#### Pyrosequencing

To quantify the abundance of *S. cerevisiae* relative to each competitor species in Competition 1, we pyrosequenced species-specific single nucleotide variants (SNVs). Previous studies have shown that pyrosequencing can accurately quantify the frequency of single nucleotide variants in pooled DNA samples (Lavebratt and Sengul 2006). To design pyrosequencing primers, we generated pair-wise sequence alignments of *ACT1* or *CYT1* between *S. cerevisiae* and each of the other species and identified at least one SNV for each pair and designed corresponding primer sets that included (1) forward and reverse primers for PCR and (2) a pyrosequencing primer (Table S2). Pyrosequencing was carried out using a PyroMark Q96 MD Automated pyrosequencer (Qiagen, Valencia, CA) following the protocol described by King and Scott-horton (2007) and the manufacturer’s directions.

To calibrate each primer set, DNA from samples containing known ratios of cells from *S. cerevisiae* and each of the other species were also pyrosequenced and used to establish standard curves using linear and polynomial regression (Figure S1). For two species (*Nakeseomyces bacillisporus* and *Tetrapisispora blattae*), we were not able to design sets of primers, and for three species (*Kazachstania lodderae*, *K. martiniae*, and *Kluyveromyces lactis*), we were not able to quantify abundance due to severely biased PCR or pyrosequencing identified by our control calibrations.

#### High-throughput sequencing

To quantify the abundance of each *S. cerevisiae* or *S. paradoxus* strain relative to each competitor species in Competition 2, we used high-throughput sequencing due to lower cost, larger sample capacity and a change in the availability of pyrosequencing. The internal transcribed sequence (ITS1) between the 5.8S and 18S ribosomal genes was amplified using 24 barcoded primers (Bokulich et al. 2012). Samples were further multiplexed using 18 indexed adaptors (Table S3). Following quantification and pooling at equal concentrations, the sample library was sequenced on a single 1x250 bp run of an Illumina MiSeq. Sample sequences were de-multiplexed allowing a single mismatch to each barcode or index followed by adaptor trimming and clipping of any low quality bases using ea-utils (Aronesty 2011). Species’ counts in each sample were obtained by blastall (v2.2.25) against 287,101 sequences from culturable species in the UNITE+INSD database (Kõljalg et al. 2013). Out of 11.6 million raw reads, 8.0 million were included in the analysis with a median of 4,346 species tags across 578 samples.

ITS1 species counts were calibrated using DNA from samples with known ratios of cells from *S. cerevisiae* or *S. paradoxus* and each of the competitor species. Standard curves were generated using linear regression (Figure S2). For three species with non-linear calibration curves, *Naumovozyma castellii*, *K. lodderae* and *Candida glabrata*, we used a modified regression:

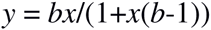

where *b* is a bias parameter, *x* is the observed frequency and *y* is the known frequency (Moskalev et al. 2011). For two species with particularly strong bias, *N. castellii* and *K. lodderae*, we also generated calibration curves for *ACT1* PCR fragments. *ACT1* PCR fragments showed a weaker bias, and so we used these fragments for all competitions between *S. cerevisiae* and either *N. castellii* or *K. lodderae*.

*Assessing dominance* – To determine the abundance of *S. cerevisiae or S. paradoxus* relative to each species, we adjusted the percentage of species-specific SNVs sequenced during pyrosequencing (Competition 1) or the number of species-specific counts from high-throughout sequencing (Competition 2) using our standard curves. If values were negative after this adjustment, we conservatively adjusted them to 0.005. To calculate the relative fitness of each competitor species, we used:

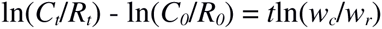

where *C*_*t*_ and *R*_*t*_ are the frequencies of the competitor and reference at generation *t* and *w* is fitness. For all competitions we assumed *t* = 14 based on changes in cell density. To identify significant growth differences between species, we either used a one-tailed paired Welch’s *t* test (Competition 1), or a mixed effect model with random effect for strain and a fixed effect for the competitor’s fitness (Competition 2). To correct for multiple comparisons, we used the method of Benjamini and Hochberg (1995) and a false discovery rate (FDR) cutoff of less than 0.01 for Competition 1 and 0.05 for Competition 2, since fewer species were competed in Competition 2.

### Intrinsic growth rate experiments

#### Growth assays

Cultures were inoculated at an optical density (OD) at 600 nm of approximately 0.25 in 1 ml of growth medium and were grown in 2 ml 96-well plates that were covered with breathable film and incubated at 30°C with shaking at 400 rpm for up to 48 hours. Cell density was measured by OD at 620 nm at 0, 4, 8, 12, 18, 24, 36, and 48 hours using an iEMS microplate reader (Thermo Lab Systems, Helsinki, Finland).

#### Analysis

The intrinsic growth rate (*r*) of each species was calculated for each time interval using the equation:

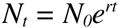

where *N*_*t*_ is final cell density, *N*_0_ is initial cell density and *t* is time in hours. The average intrinsic growth rate of each species across all sampling intervals for a given medium was then used to evaluate the effect of a treatment (*i.e*., low-pH or a nutrient supplement) on the growth of each species, (*Δr*_*Treatment*_ = *r*_*Treatment*_ – *r*_*Control*_), or the effect of the species in a given environment (*Δr*_*Species*_ = *r*_*non-S. cerevisiae*_ *r*_*S. cerevisiae*_). While the average growth rate does not distinguish between differences in lag phase, exponential growth rate and carrying capacity, we used it because it had a smaller error among replicates.

For two sets of experiments, we only used OD up to and including 24 hours in our analysis: ethanol tolerance experiments and supernatant experiments. For ethanol tolerance experiments we observed flocculation that increased the variability of OD measurements beginning at 36 hours. For inhibitor compound experiments we observed that un-inoculated control samples registered noticeable effects on OD measurements beginning at 36 hours.

Ethanol tolerance among species was measured by the ethanol concentration that inhibited growth by 50% (IC_50_). IC_50_ estimates were obtained by fitting dose response curves using a three-parameter Weibull function in R (R Development Core Team 2013) using the ‘drc’ package (Ritz and Streibig 2005). Statistical comparisons between the estimated IC_50_ for *S. cerevisiae* and each species were made using the ‘comped’ function (Ritz and Streibig 2005; R Development Core Team 2013) followed by the Altman and Bland method to calculate *P* values from confidence intervals (Altman and Bland 2011). To correct for multiple comparisons, we used the method of Benjamini and Hochberg (1995) and a FDR cutoff of less than 0.01 for significance.

## Results

Ecological dominance of yeast species in grape juice evolved more recently than the fermentative lifestyle

To determine when ecological dominance of high-sugar environments evolved, we grew representative strains of multiple non-WGD and WGD yeast species in two high-sugar environments: Chardonnay grape juice (Grape) and a high-sugar rich medium (HS). If dominance in high-sugar environments evolved along with the evolution of fermentation in the presence of oxygen, we expect *S. cerevisiae* will consistently exhibit higher relative fitness than all non-WGD species but not WGD species in both high-sugar environments. Grape was chosen to represent a natural high-sugar environment, and HS was chosen to replicate the sugar concentration typical of the grape juice environment while limiting the potential influence of nutrient content and low-pH. Since the ability to dominate is inherently a relative trait, we assessed the growth of each non-WGD and WGD species relative to a representative *S. cerevisiae* strain isolated from an oak tree (YPS163) using co-cultures. As a control, we also grew a *S. cerevisiae* strain isolated from a vineyard (I14) in co-culture with our reference *S. cerevisiae* strain. If the relative abundance of *S. cerevisiae* was significantly higher at the end compared to the start of the experiment, indicating a higher relative fitness, it was considered “dominant”.

We find that *S. cerevisiae* dominates nearly all non-WGD and WGD yeast species in Grape and HS co-cultures (Figure 2A and 2B). In both Grape and HS, *S. cerevisiae* increased in abundance relative to 10/12 yeast species (FDR < 0.01, Table S4). In the majority of these co-cultures, *S. cerevisiae* was greater than 90% of the population at 48 hours. Notably, *S. cerevisiae* remained a significant proportion of the population even when it did not dominate. These data show that *S. cerevisiae* is able to dominate in multiple high-sugar environments, and they suggest that the ability to dominate high-sugar environments arose recently in yeast evolution.

**Figure. 2.**
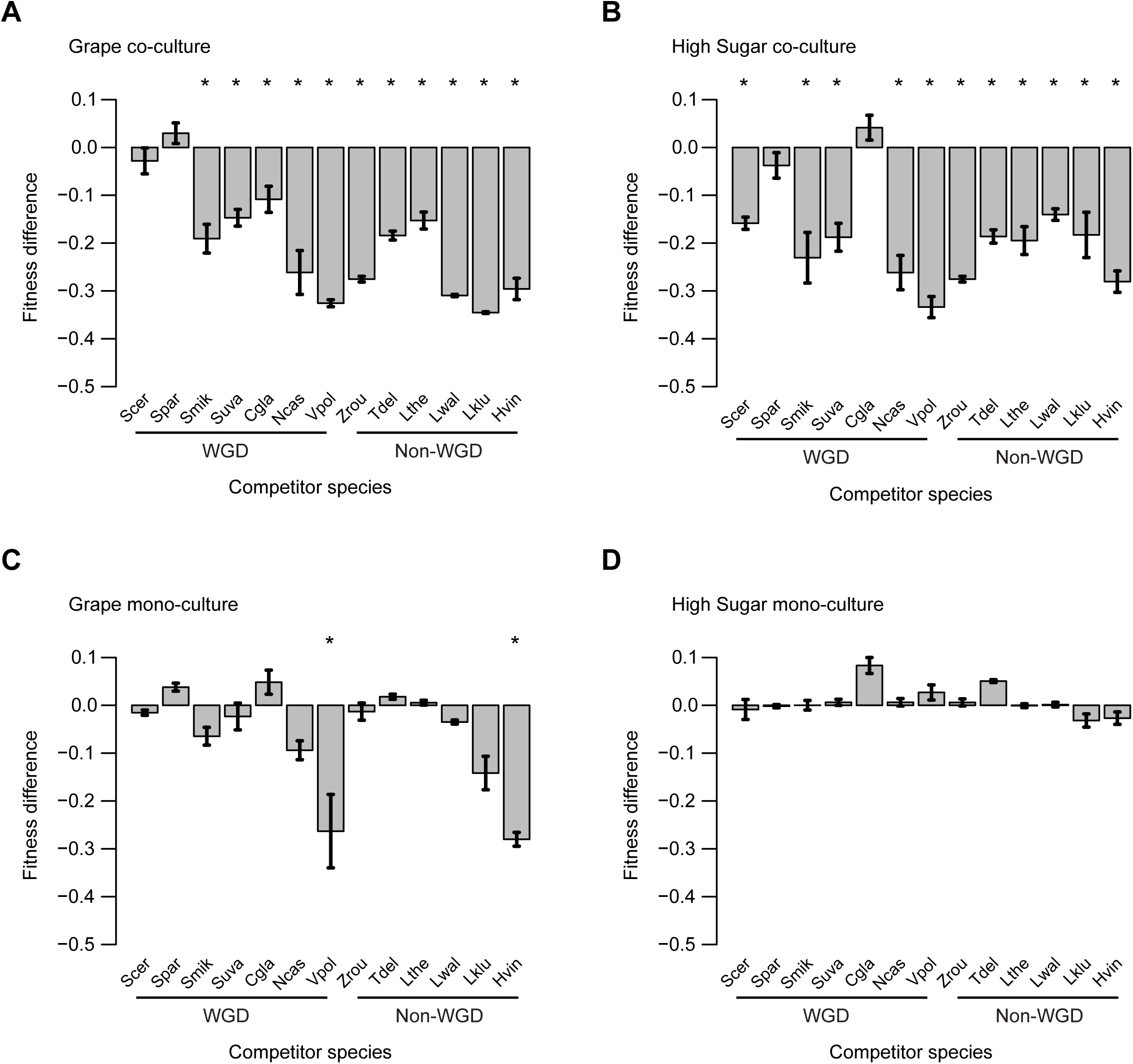
Fitness differences of *S. cerevisiae* relative to WGD and non-WGD yeast species after co-culture or mono-culture in two high-sugar environments. The fitness of each species in Grape (A) and HS (B) co-cultures and Grape (C) and HS (D) mono-cultures. Bars and whiskers represent the mean and standard error (n = 3) of the difference between the competitor fitness (*w*_*c*_) and the *S. cerevisiae* (YPS163) reference fitness (*w*_*r*_), which is set to one. Species labels are Scer (*S. cerevisiae*), Spar (*S. paradoxus*), Smik (*S. mikatae*), Suva (*S. uvarum*), Cgla (*C. glabrata*), Ncas (*N. castellii*), Vpol (*V. polyspora*), Tdel (*T. delbruekii*), Lthe (*L. thermotolerans*), Lwal (*L. waltii*), Lklu (*L. kluyveri*), Hvin (*H. vineae*) and WGD and non-WGD species are indicated. Fitness significantly different from the reference is shown for FDR < 0.01 (*).

In support of a more recent evolution of ecological success in high-sugar environments, *S. paradoxus* is the only species that persists along with *S. cerevisiae* in Grape and HS co-cultures. Two other strains, *S. cerevisiae* (I14) and *C. glabrata,* also competed well with our *S. cerevisiae* reference. However, their persistence depended upon the environment: *S. cerevisiae* dominated *C. glabrata* in Grape (FDR = 0.0072) but not in HS, whereas it dominated I14 in HS (FDR = 0.0010) but not in Grape. Thus, *S. paradoxus* was the only species able to compete well with *S. cerevisiae* in both high-sugar environments.

One explanation for *S. cerevisiae*’s dominance in our Grape and HS co-cultures is that it has a greater carrying capacity than other species in these environments, even when they are grown individually. As a control for our co-culture experiments, we also measured the density of each species grown in mono-culture by mixing it with a *S. cerevisiae* mono-culture after 48 hours and quantifying the proportion of each species.

*S. cerevisiae* has a carrying capacity similar to the majority of yeast species in Grape and HS (Figure 2C and 2D). In Grape, the abundance of *S. cerevisiae* was significantly greater than only 2/12 species after 48 hours of mono-culture (Figure 2C, FDR < 0.01). Species that obtained significantly lower carrying capacities included the non-WGD species *Hanseniaspora vineae* and the WGD species *Vanderwaltozyma polyspora*, which were 1% and 3% of *S. cerevisiae*’s abundance after 48 hours in mono-culture. The relative population size of *S. cerevisiae* was also not significantly greater than 12/12 species tested in HS. These data imply that *S. cerevisiae*’s dominance in Grape and HS co-cultures is not due to differences between species in their individual carrying capacities.

Dominance of high-sugar environments appears to be a trait shared by *S. cerevisiae* and *S. paradoxus*. However, we only used one representative strain of *S. cerevisiae* and did not compete *S. paradoxus* with other species. To test whether *S. cerevisiae*’s dominance is a property of the strain used or the species we repeated a subset of the Grape and HS competitions using two additional strains of *S. cerevisiae*: a soil isolate (I14) from Italy that is closely related to other European strains (Fay and Benavides 2005) and a tree isolate from China that is diverged from both the European strain and our reference strain YPS163 from North America (Wang et al. 2012). The fitness of the three *S. cerevisiae* strains together in relation to each competitor species indicates that *S. cerevisiae*: dominates *S. mikatae*, *S. uvarum*, *N. castelli* and *K. lodderae* in both Grape and HS, dominates *C. glabrata* in Grape but not HS, and has no fitness differences in comparison to *S. paradoxus* (FDR < 0.05, Figure 3A and 3B). These results confirm our previous results based on a single *S. cerevisiae* reference strain and add *K. lodderae* to the list of species dominated by *S. cerevisiae*.

**Figure. 3.**
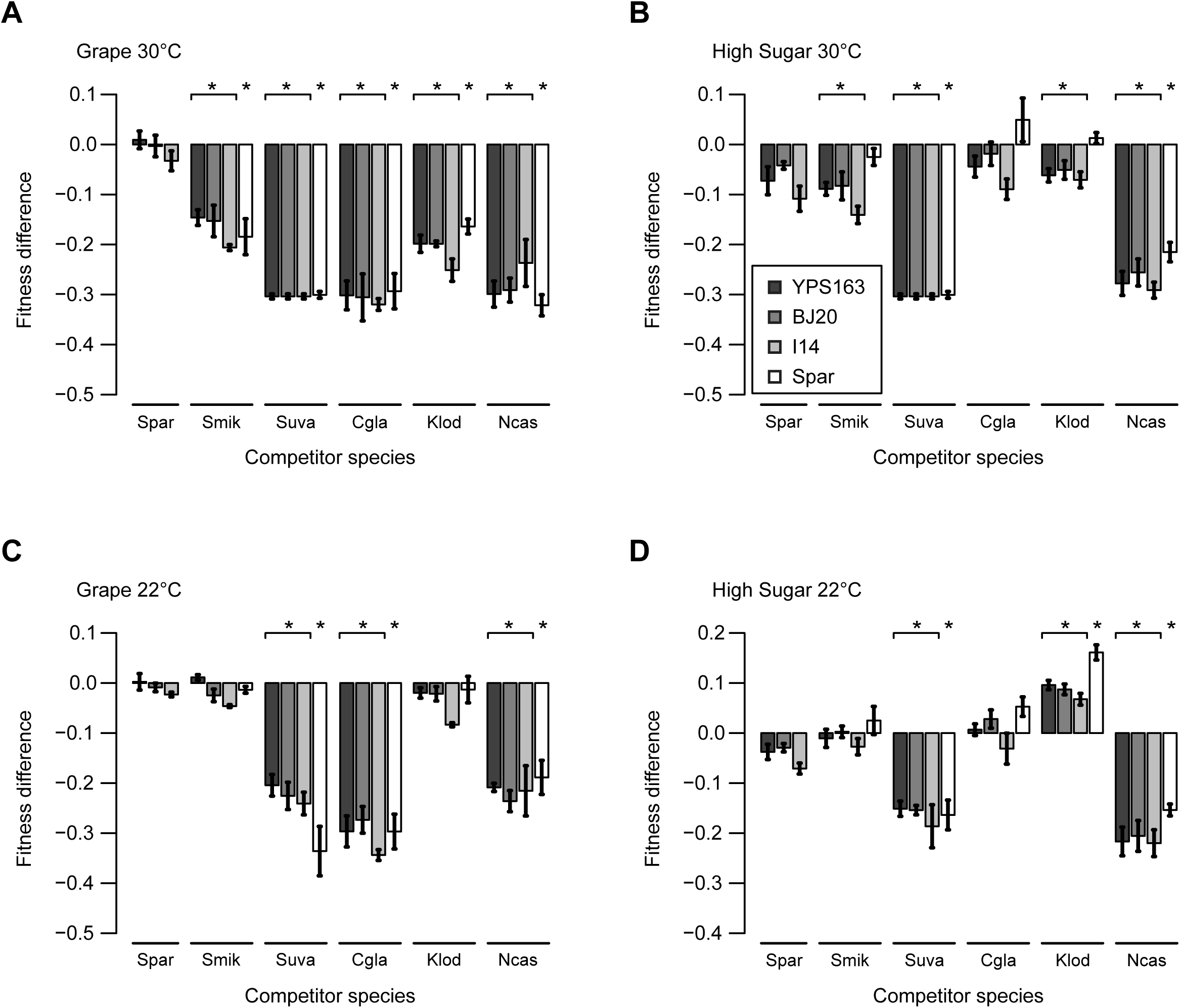
Fitness differences depend on temperature and growth medium. Fitness differences between each competitor species and a reference strain are shown for competitions at 30°C in Grape (A) or HS (B) and at 22°C in Grape (C) or HS (D) medium. Reference strains are three *S. cerevisiae* strains (YPS163, BJ20 and I14) and *S. paradoxus* (Spar). Bars and whiskers represent the mean and its standard error (n = 3). Significant differences in fitness are shown for FDR < 0.05 (*). Species abbreviations are the same as in Figure 2.

Using *S. paradoxus* as the reference species we find that *S. paradoxus* dominates the same yeast species as *S. cerevisiae* in Grape (Figure 3A). In HS, *S. paradoxus* dominates fewer species than *S. cerevisiae*; it does not dominate *S. mikatae* and *K. lodderae* (Figure 3B). Thus, the results of our competition experiments with *S. paradoxus* show that dominance in Grape is shared by both *S. cerevisiae* and *S. paradoxus* but that the ability of *S. paradoxus* to dominate in HS is not as strong as that of *S. cerevisiae*.

Temperature affects ecological dominance of certain yeast species in high-sugar environments

The *Saccharomyces* species have differentiated in both their optimal and maximum growth temperature (Belloch et al. 2008; Gonçalves et al. 2011; Salvadó et al. 2011b; Kurtzman et al. 2011). Because *S. cerevisiae*’s optimal growth temperature (32°C), is higher than that of *S. paradoxus* (30°C), *S. mikatae* (29°C) and *S. uvarum* (26°C) (Salvadó et al. 2011b), its dominance of high-sugar environments could depend on the high temperature (30°C) of our initial competition experiment. To examine this possibility, we competed a subset of species at a lower temperature (22°C).

Similar to the results of our competition experiments performed at high temperature, at 22°C both *S. cerevisiae* and *S. paradoxus* dominated *S. uvarum* and *N. castellii* in Grape and HS, and dominated *C. glabrata* in Grape but not HS (Figure 3C and 3D). However, at the lower temperature neither *S. cerevisiae* nor *S. paradoxus* dominated *S. mikatae* or *K. lodderae* in Grape or HS, and *K. lodderae* dominated *S. cerevisiae* and *S. paradoxus* in HS. Thus, ecological dominance of certain yeast species in high-sugar environments depends on temperature, which implicates the evolution of thermal tolerance among the *Saccharomyces* species in the evolution of ecological dominance.

*S. cerevisiae* has a distinct competitive advantage in grape juice

Our finding that many WGD yeast species compete poorly with *S. cerevisiae* in high-sugar environments suggests that the fermentative lifestyle is not sufficient to confer ecological success in these environments. Since the majority of the species we tested are capable of achieving similar carrying capacities to *S. cerevisiae* in these environments when grown individually, *S. cerevisie*’s dominance in our Grape and HS co-cultures must be related to either differences in intrinsic growth rates or interference competition. To investigate these two modes of ecological dominance, we measured the intrinsic growth rate of each species in mono-culture. If *S. cerevisiae* does not exhibit a greater intrinsic growth rate than other species, then its ability to dominate in these environments can be attributed to interference competition.

*S. cerevisiae* has a greater intrinsic growth rate than nearly all yeast species in the grape juice environment (Figure 4). Compared to *S. cerevisiae,* 16/17 species exhibited a significantly lower intrinsic growth rate in Grape (FDR < 0.01, Table S5). The one notable exception to this pattern was *S. paradoxus*, which had a lower growth rate but did not meet our cutoff for significance (FDR = 0.028). In stark contrast to our finding in Grape, when we compared the intrinsic growth rate of *S. cerevisiae* to the intrinsic growth rate of each of the other species in HS, we did not observe any significant difference for 17/17 species (Figure 4B). These results suggest different or multiple mechanisms contribute to the dominance of *S. cerevisiae* in high-sugar environments. Furthermore, they support the recent evolution of traits required for ecological success in the grape juice environment. In the following sections we examine factors that may contribute to the dominance of *S. cerevisiae* in both HS and Grape.

**Figure. 4.**
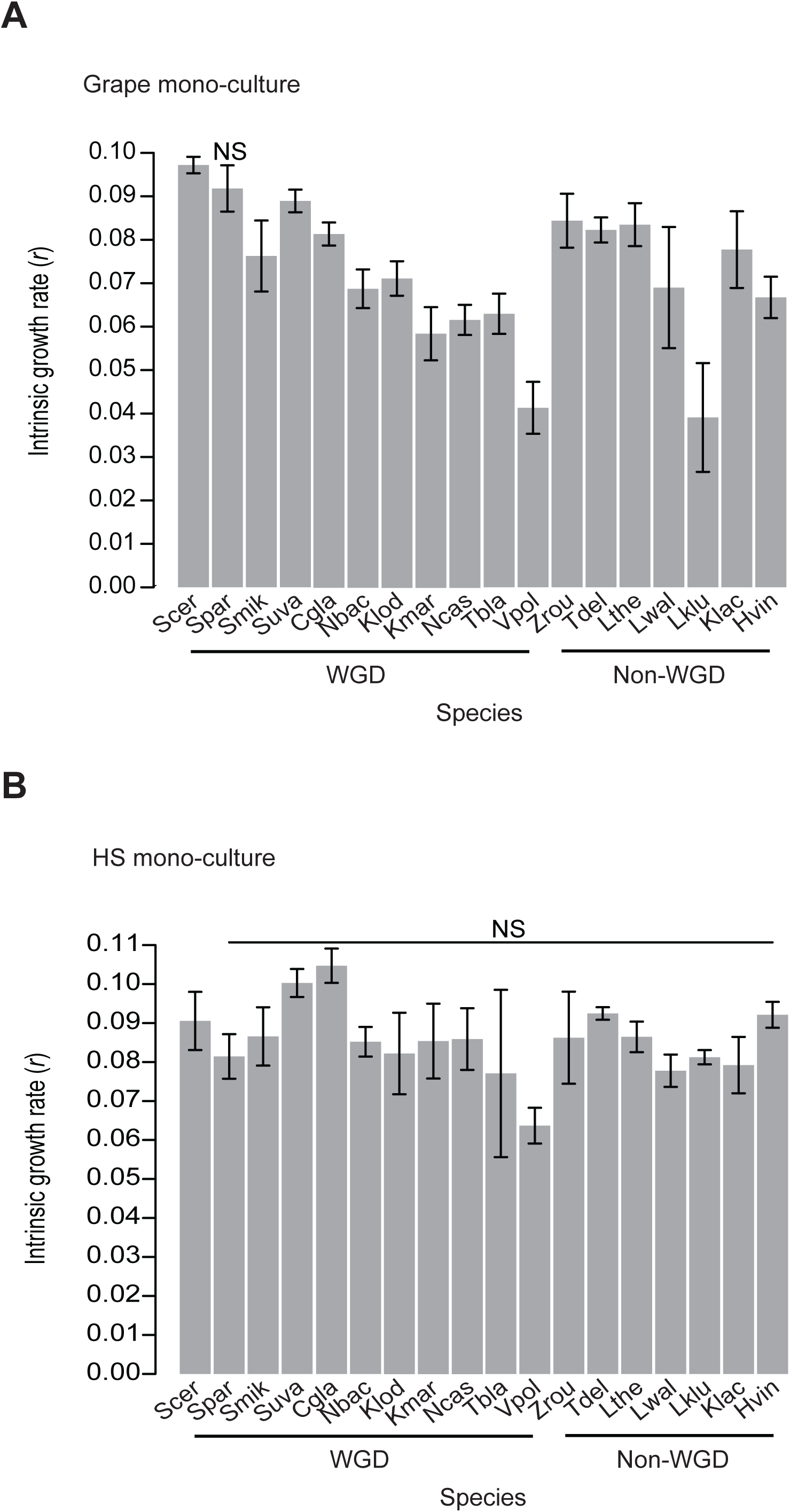
Intrinsic growth rate differences in high-sugar environments. The intrinsic growth rate of each species in Grape (A) and HS (B). WGD and non-WGD species are indicated. Bars and whiskers represent the mean and standard deviation of the growth rate (n = 3). Species that did not differ significantly from *S. cerevisiae* at an FDR cutoff of 0.01 are indicated (NS).

### Evolution of ethanol tolerance and its potential role in interference competition

*S. cerevisiae*’s ability to produce and tolerate ethanol is one way in which it may dominate other species in high-sugar environments. Although previous studies showed that *S. cerevisiae* tolerates higher ethanol concentrations than many yeast species, they only included 4/17 of the species used in this study (Pina et al. 2004; Belloch et al. 2008; Arroyo-López et al. 2010; Salvadó et al. 2011a). To examine the potential impact of ethanol on the growth of each species, we measured the intrinsic growth rate of each species in YPD supplemented with ethanol at concentrations ranging from 0-10% and calculated the ethanol concentration that inhibited growth rate by 50% (IC_50_) for each species by fitting dose-response curves to the growth rate (see Materials and Methods).

*S. cerevisiae* had an IC_50_ greater than 15/17 yeast species (Figure 5 and Table S6). The two exceptions to this pattern were *S. cerevisiae*’s closest relative, *S. paradoxus* (FDR = 0.0502) and *C. glabrata* (FDR = 0.0320). *C. glabrata* grew as well as *S. cerevisiae* at moderate ethanol concentrations, and it grew better than *S. cerevisiae* at low ethanol concentrations (Figure S3 and Table S6). However, most of *S. cerevisiae*’s growth advantage occurred at ethanol concentrations at or above 4% (Figure S3 and Table S6). Thus, while all species tolerate low concentrations of ethanol (< 4%), *S. cerevisiae* exhibits a growth advantage compared to most species at high-ethanol concentrations.

**Figure. 5.**
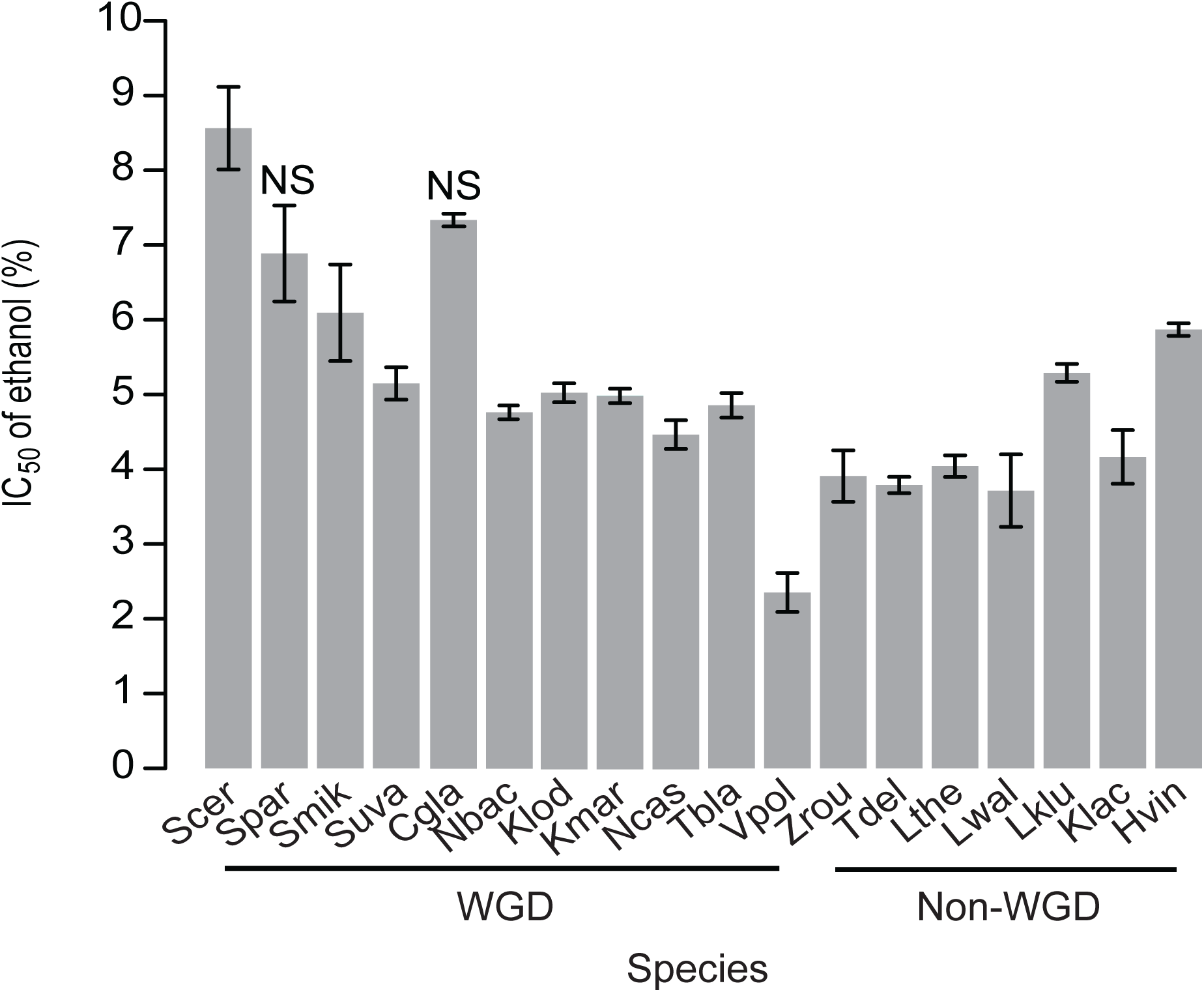
Species differences in ethanol tolerance. Mean (bars) and standard error (whiskers) of the concentration of ethanol (%) that inhibits growth by 50% (IC_50_) of each yeast species (n = 3). WGD and non-WGD species are indicated. Species with an IC_50_ that did not differ significantly from *S. cerevisiae* at an FDR cutoff of 0.01 are indicated (NS).

### No evidence for interference competition mediated by other toxic metabolites

Previous studies found that *S. cerevisiae* produced toxic metabolites other than ethanol that inhibit the growth of competitor species (van Vuuren and Jacobs 1992; Magliani et al. 1997; Musmanno et al. 1999; Pérez-Nevado et al. 2006; Albergaria et al. 2010; Rodríguez-Cousiño et al. 2011; Branco et al. 2014). However, other studies either did not find any evidence that *S. cerevisiae* produced an inhibitory compound (Torija et al. 2001; Nissen et al. 2003; Arroyo-López et al. 2011) or found that the ability to produce killer toxins varied among *S. cerevisiae* strains (Gutiérrez et al. 2001; Sangorrín et al. 2007; Maqueda et al. 2012). To determine whether the *S. cerevisiae* strain we used during our assays produces an inhibitor compound, we grew each species in the supernatant obtained from YPD mono-cultures and co-cultures with *S. cerevisiae.* We chose YPD, which contains 2% dextrose, because ethanol concentrations should not attain inhibitory concentrations during growth. In no instance did the supernatant inhibit the subsequent growth of each species (Figure S4 and Table S7).

Low-pH and nutrient limitations contribute to *S. cerevisiae*’s intrinsic growth rate advantage in grape juice

Grape juice differs from high-sugar rich medium in that it has a lower pH (pH = 3.7 vs pH = 6.7) and reduced levels of nutrients, most notably yeast assimilable nitrogen (Henschke and Jiranek 1993). To determine whether *S. cerevisiae*’s higher intrinsic grow rate in grape juice is related to pH or nutrient deficiencies we measured the effects of altered pH of HS and nutrient content of Grape for each species.

To test the effect of pH on the intrinsic growth rate of each species, we grew each species in low-pH HS, HS adjusted to the same acidity level as our Grape medium. As a control, we compared each species’ growth in low-pH HS to its growth in HS. If *S. cerevisiae*’s intrinsic growth rate is greater than other species in Grape due to low-pH, then *S. cerevisiae* should also exhibit a higher intrinsic growth rate than other species in low-pH HS.

*S. cerevisiae* has an intrinsic growth rate advantage in low-pH HS (Figure 6).

**Figure. 6.**
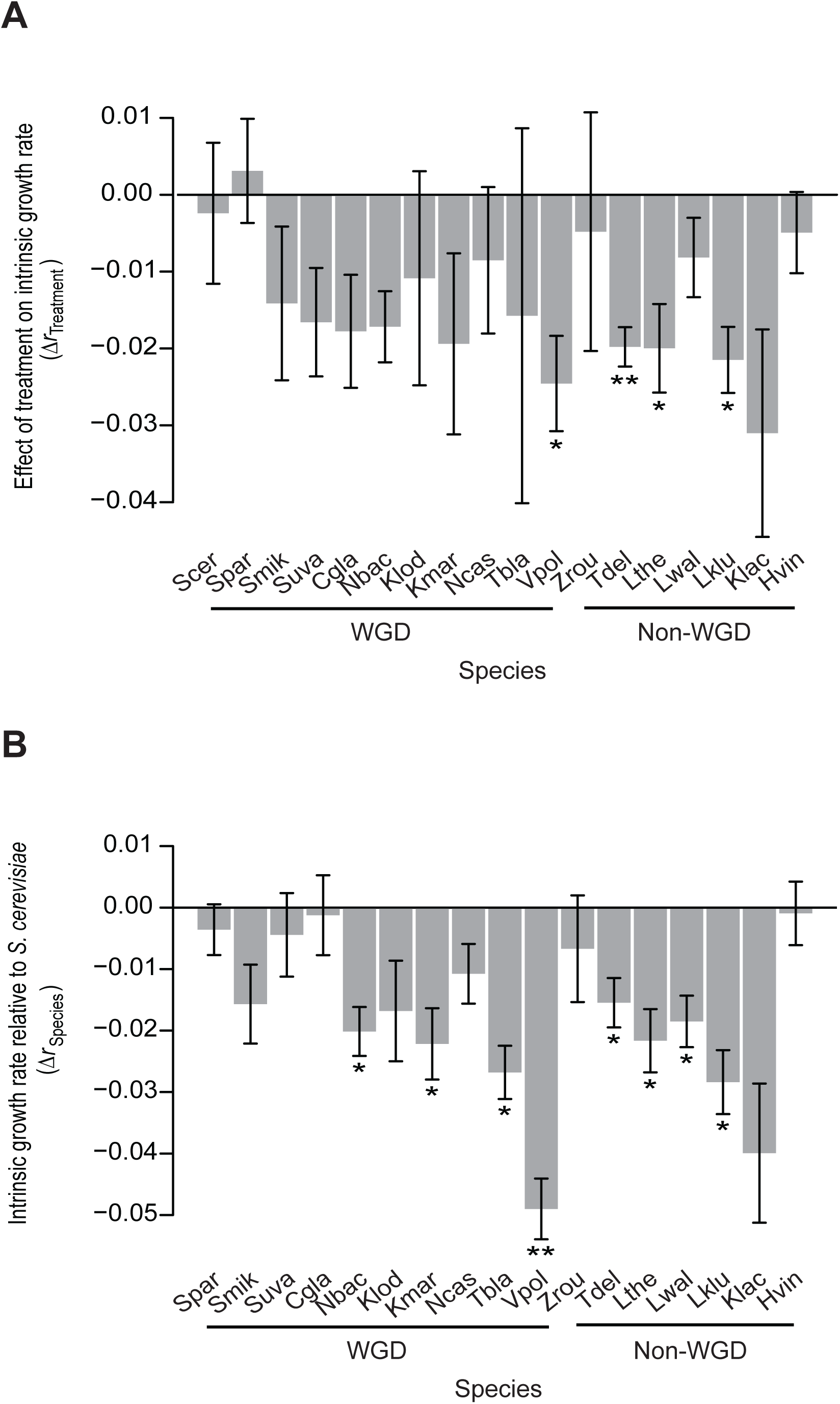
Intrinsic growth differences in response to low-pH. (A) The effect of low-pH treatment on the intrinsic growth rate (*r*) of each species in HS (*Δr*_*Treatment*_ = *r*_*Treatment*_ – *r*_*HS*_), and (B) the difference in the intrinsic growth rate between *S. cerevisiae* and each species in low-pH HS (*Δr*_*Species*_ = *r*_*non-S. cerevisiae*_ – *r*_*S. cerevisiae*_). WGD and non-WGD species are indicated. Whiskers for each bar show 95% confidence intervals (n = 3). Significant differences in the growth rate of each species with or without low-pH and differences between *S. cerevisiae* and each species in low-pH are shown for FDR < 0.01 (*) and FDR < 0.001 (**).

When grown in low-pH HS, 4/18 species exhibited a significantly lower intrinsic growth rate when compared to growth in HS (Figure 6A and Table S8). Notably, only three species, including *S. cerevisiae* were not affected by low-pH at a nominal level of significance (*P* < 0.05) compared to an FDR cutoff of 0.01. Additionally, when we compared the intrinsic growth rate of *S. cerevisiae* in low-pH HS to each of the other species in this environment, *S. cerevisiae*’s intrinsic growth rate was greater than 8/17 species (Figure 6B), compared to 0/17 species observed in HS (Figure 4B).

To test the effect of nutrient deficiency on the intrinsic growth rate of each species, we grew each species in Grape supplemented with one of several different nutrient sources that varied in complexity: YP, CM, NB, AA, and AS. YP is the rich nutritive base of the HS environment, CM contains vitamins and minerals, amino acids, and a single good nitrogen source, and NB, AA, and AS are the vitamins and minerals, amino acids, and nitrogen source (ammonium sulfate) components of CM, respectively. As a control, we compared each species’ growth in Grape with a nutrient supplement to its growth in Grape without the nutrient supplement. If nutrient limitations contribute to intrinsic growth rate differences between species, then nutrient supplements in Grape should increase each species’ growth rate and reduce or eliminate intrinsic growth rate differences between species.

Most yeast species are nutrient limited in grape juice. Of the 18 species we assayed, 12 exhibited a significant increase in intrinsic growth rate with the addition of one or more nutrient supplements, including *S. cerevisiae* (Figure 7, Figure S5 A-D, and Table S9). However, which nutrients elicited a significant increase in growth varied by species. For example, *S. uvarum* was positively affected by the addition of YP and NB, whereas *C. glabrata* was positively affected by YP, CM and AA. Overall, YP positively affected the intrinsic growth rate of the most species (11), followed by NB (7), CM (6), and AA (3). However, none of the species we assayed grew significantly better with the addition of AS, a good nitrogen source.

**Figure. 7.**
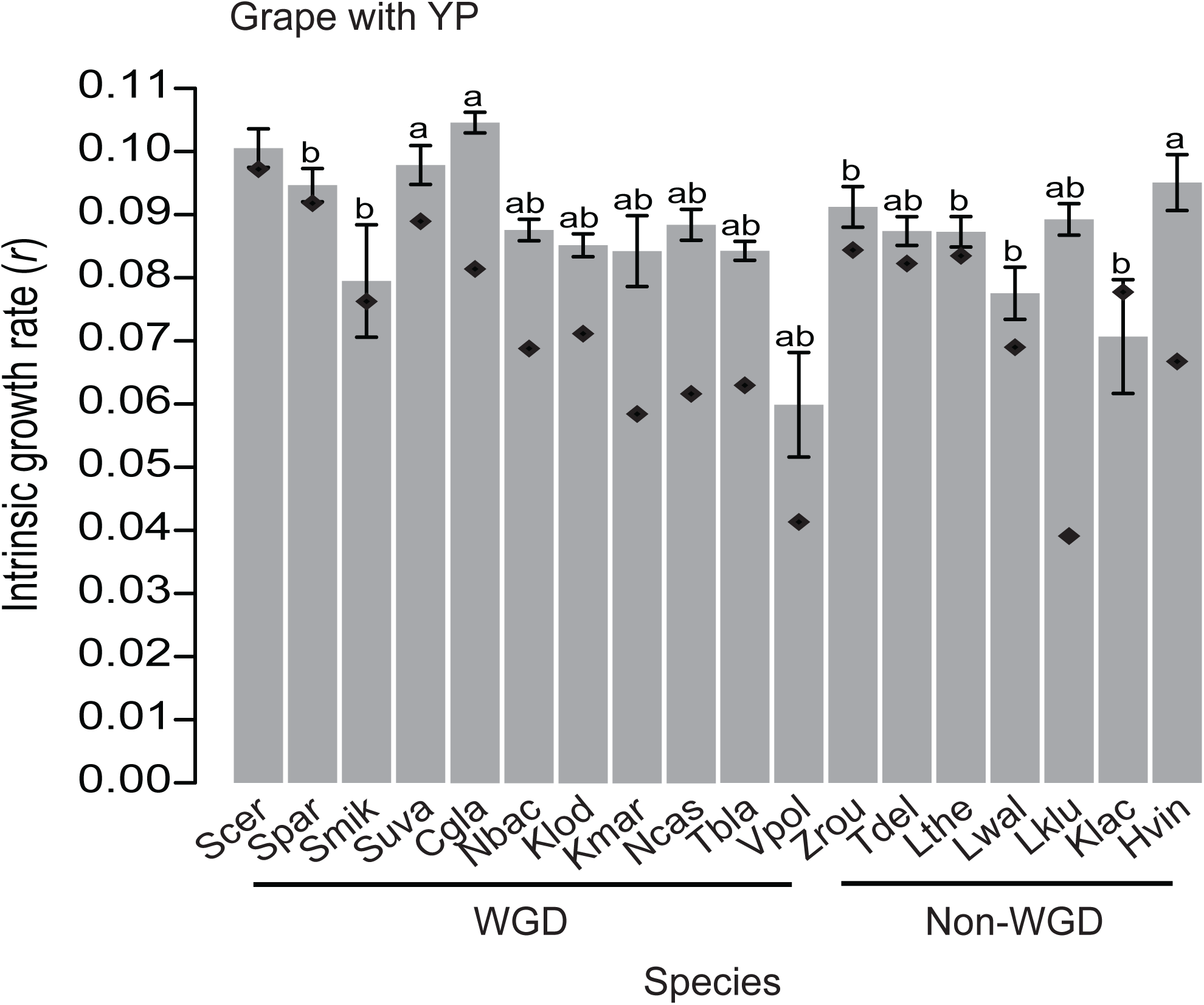
Intrinsic growth rate in Grape supplemented with nutrients. The mean intrinsic growth rate of each species in Grape supplemented with nutrients (YP). WGD and non-WGD species are indicated. Bars and whiskers represent the mean and standard deviation of the growth rate (n = 3). Diamonds represent the mean growth rate in Grape without YP. Significant differences in the growth rate of each species with or without YP (a) and differences between *S. cerevisiae* and each species in YP (b) are labeled above each bar for FDR < 0.01.

Nutrient supplements eliminate the intrinsic growth rate differences between *S. cerevisiae* and nearly all other yeast species. Of the 17 species that grew significantly slower than *S. cerevisiae* in Grape (Figure 4A), only two species, *N. castellii* and *V. polyspora*, grow significantly slower than *S. cerevisiae* in spite of all of the nutrient supplements used in our experiments (Table S9). Overall, Grape supplemented with CM had the fewest number of species that still grew significantly slower than *S. cerevisiae* (4), followed by Grape supplemented with AS (7), NB (12), YP (14), and AA (16).

## Discussion

The diversion of more sugar to fermentation than respiration in the presence of oxygen, i.e. the fermentative lifestyle, provides many yeast species the opportunity to exploit novel environments and ecological strategies. One such species, *S. cerevisiae*, consistently dominates wine fermentations and has become widely used to ferment beer, bread and wine. In this study, we investigated when *S. cerevisiae’s* ability to dominate high-sugar environments evolved and whether its dominance is a simple consequence of the fermentative lifestyle. We find that dominance evolved much more recently than the evolution of the fermentative lifestyle and for certain species depends on the temperature and growth medium. Our results suggest that *S. cerevisiae*’s frequently observed dominance of grape juice fermentations is mediated by the evolution of multiple traits that build on an ancient change in metabolism.

### Evolution of dominance in relation to the fermentative lifestyle

Our study indicates that dominance in high-sugar environments evolved much more recently than the WGD and the transition to the fermentative lifestyle (Hagman et al. 2013). While previous studies showed that *S. cerevisiae* dominates multiple other non-WGD species (Holm Hansen et al. 2001; Fleet 2003, 2008; Pérez-Nevado et al. 2006; Goddard 2008), our findings demonstrate that *S. cerevisiae* and its closest relative, *S. paradoxus* also dominate most other WGD yeast species in high-sugar environments and that the temperature and growth medium are also important. The ability of *S. cerevisiae* and *S. paradoxus* to dominate representatives of multiple taxonomically diverse species suggests that dominance arose quite recently. However, the change in dominance relationships across environments implies that: i) multiple traits underlie dominance, ii) these traits have changed on lineages other than that leading to the ancestor of *S. cerevisiae* and *S. paradoxus*, and iii) dominance cannot be ascribed to a single lineage.

Multiple, distantly-related WGD lineages grow well in nutrient rich, high-sugar environments, but few species grow as well as *S. cerevisiae* in grape juice. While *S. cerevisiae* and *S. paradoxus* dominate most WGD yeast species in HS medium, they do not dominate *C. glabrata* in this environment, and depending upon the temperature, they also lose some or all of their competitive advantage over *S. mikatae* and *K. lodderae*. Remarkably, *K. lodderae*’s higher relative fitness in HS compared to Grape contributes to its unique ability to dominate *S. cerevisiae* and *S. paradoxus* in HS at low-temperature. One explanation for the exceptional growth observed for these species is that the ability to compete in nutrient rich, high-sugar environment evolved independently along the lineages that gave rise to *K. lodderae*, *C. glabrata,* and the *Saccharomyces* species. Alternatively, the ability to compete well in nutrient rich, high-sugar environments may have evolved early during the transition to the fermentative lifestyle, followed by multiple, independent losses. However, our findings suggest that the more parsimonious explanation is that the ability to dominate evolved much more recently than the transition to the fermentative lifestyle.

Evolution of temperature preferences affects the dominance of certain species. Among the *Saccharomyces* species, *S. cerevisiae*, *S. paradoxus* and *S. mikatae* can grow at 37°C, whereas *S. arboricolus*, *S. uvarum* and *S. kudriavzevii* cannot and the latter two are considered cryophilic (Belloch et al. 2008; Gonçalves et al. 2011; Salvadó et al. 2011b). Outside the *Saccharomyces* species, *K. lodderae*, *K. martiniae*, *T. blattae*, *L. waltii* and *H. vineae* cannot grow at 37°C (Kurtzman et al. 2011), indicating that growth at high temperature has been gained and/or lost multiple times. Given the species tree (Figure 1), we attribute *S. cerevisiae*’s and *S. paradoxus*’ dominance of *S. mikatae* at 30°C but not 22°C to evolution of thermal tolerance. Similarly, *S. cerevisiae*’s dominance of *K. lodderae* in Grape at 30°C but not 22°C may also depend on the evolution of thermal tolerance. Although we did not specifically assay dominance below 22°C, the outcome of competitions performed at even cooler temperatures may change. *S. uvarum* dominates some wines at low temperatures (Torriani et al. 1999; Naumov et al. 2000; Sipiczk et al. 2001; Rementeria et al. 2003; Demuyter et al. 2004), and *S. kudriavzevii* does not dominate but competes better with *S. cerevisiae* at low temperatures (Arroyo-López et al. 2011). However, *S. cerevisiae*’s dominance of *S. uvarum* may not be a simple consequence of thermal tolerance; *S. uvarum*’s growth rate in grape juice increased with supplementation of rich medium (YP) to a rate equivalent to that of *S. cerevisiae*. Furthermore, we did not observe a significant intrinsic growth rate difference between these species in high-sugar rich medium when grown at 30°C (Figure 4).

The thermal differentiation of *Saccharomyces* species raises the possibility that the ability to dominate high-sugar environments evolved prior to their differentiation. Dominance of high-sugar environments may have evolved progressively such that WGD yeast species can outcompete non-WGD yeast species even though they lose to *S. cerevisiae*. However, more pair-wise competitions between other non-Saccharomyces WGD species and non-WGD species are needed to test that hypothesis. While the evolution of dominance cannot be pinned to any particular lineage, we see no clear progression to higher ethanol tolerance among WGD yeast species or evidence for higher intrinsic growth rate in grape juice outside of the *Saccharomyces* species. Thus, in addition to thermal differentiation, other traits likely to contribute to dominance evolved among the *Saccharomyces* species.

One limitation of our ability to make inferences about the evolution of dominance and how the environment influences it is that most species were only represented by a single strain. While the fitness of three diverse strains of *S. cerevisiae* are generally quite similar to one another and to *S. paradoxus*, competitor strains may not always be representative of the species. This possibility places some limits on the interpretation of dominance relationships specific to a single species, e.g. *C. glabrata* and *K. lodderae*, but is unlikely to explain overall patterns of dominance.

### Multiple mechanisms of ecological dominance

*S. cerevisiae*’s dominance of high-sugar environments cannot be explained by a single mechanism. In addition to thermal tolerance discussed above, we also find evidence for ethanol tolerance and a high intrinsic growth rate in grape juice. Interference competition through the production of ethanol provides one explanation for *S. cerevisiae*’s dominance of high-sugar rich medium. Consistent with previous studies (Pina et al. 2004; Belloch et al. 2008; Arroyo-López et al. 2010; Salvadó et al. 2011a), we found that *S. cerevisiae* exhibits greater ethanol tolerance than most species. In support of the role of ethanol tolerance in dominance, *S. paradoxus* and *C. glabrata* exhibited ethanol tolerance similar to *S. cerevisiae* and were the only two species that were not dominated by *S. cerevisiae* in high-sugar rich medium at high temperature.

However, previous studies showed that oxygen, cell density and an inhibitory peptide affect *S. cerevisiae*’s dominance of various non-WGD species (Holm Hansen et al. 2001; Nissen et al. 2003, 2004; Pérez-Nevado et al. 2006; Albergaria et al. 2010; Branco et al. 2014). These studies excluded the effects of ethanol because non-WGD species initiated cell death by some other mechanism before ethanol reached inhibitory concentrations. While we only measured competitions with two of the species used in earlier studies, *T. delbrueckii* and *L. thermotolerans*, we cannot exclude the possibility that these species were dominated for reasons other than ethanol inhibition. One difference between our experiments and those of prior studies is that they were carried out with low or no agitation, whereas we performed our competitions under high agitation (400 rpm). Agitation is expected to increase dissolved oxygen and might eliminate cell density and confinement effects (Nissen et al. 2003, 2004; Arneborg et al. 2005).

Multiple lines of evidence suggest that *S. cerevisiae*’s dominance in grape juice is influenced by its high intrinsic growth rate in this environment. *S. cerevisiae* exhibited the highest rate of growth in grape juice, significantly higher than all species except *S. paradoxus*. The absence of any difference in growth rate in high-sugar rich medium implies that *S. cerevisiae*’s intrinsic growth advantage in grape juice is specific to grape juice or similar environments. Furthermore, lowering the pH of high-sugar rich medium did not affect *S. cerevisiae* but affected the growth of other species, and supplementation of nutrients to grape juice increased the growth of many species but had little to no effect on *S. cerevisiae*. Notably, *S. uvarum*, *C. glabrata* and *H. vineae* grew as well as *S. cerevisiae* in low-pH medium and in grape juice supplemented with rich nutrients (YP), indicating that low nutrients alone may explain their slow growth in grape juice.

The relative importance of temperature, intrinsic growth rate and ethanol inhibition to *S. cerevisiae*’s dominance of grape juice is uncertain since their effects are difficult to disentangle from one another. In support of temperature, fermentation is exothermic and can increase the temperature of wine must by as much as 10°C (Boulton 1979; Goddard 2008). However, we found that nutrient supplementation increased most species’ intrinsic growth rate in grape juice at 30°C. Ethanol inhibition is not likely to be important until the later stages of fermentation because most species were not significantly inhibited by ethanol concentrations below 5%, similar to previous reports (Goddard 2008; Salvadó et al. 2011a). As such, we favor intrinsic growth rate in grape juice as a driver of dominance as it likely acts earlier and throughout the competition.

Interactions between factors may also contribute to *S. cerevisiae*’s dominance. Ethanol and high-temperature act synergistically to decrease growth due to their overlapping effects on lipid membrane integrity (Piper 1995). Lipid membrane integrity importantly affects proton (H+) transport across the cell membrane, and the combined effects of ethanol and high-temperature increase the lipid membrane’s H+ permeability (Madeira et al. 2010). Increased H+ permeability can also result in reduced intracellular pH, particularly in acidic environments such as grape juice. While we did not measure any interaction effects, Goddard (2008) found interactions between the effects of temperature, ethanol and media, including grape juice, on the growth rate of *Saccharomyces* versus non-*Saccharomyces* species.

### Ecology of high-sugar environments

The fermentative lifestyle is hypothesized to coincide with the evolution of flowering plants due to the abundance of diverse high-sugar environments (Wolfe and Shields 1997; Piskur and Langkjaer 2004; Thomson et al. 2005; Conant and Wolfe 2007). While the ecology of fermentative species is not well known, many have been isolated from insects and may be transported to high-sugar environments (Kurtzman et al. 2011). However, the recent evolution of traits that contribute to *S. cerevisiae*’s dominance of high-sugar environments is perplexing. Although both *S. cerevisiae* and *S. paradoxus* can be found in vineyards (Redzepović et al. 2002; Hyma and Fay 2013), these and other *Saccharomyces* species are commonly associated with tree bark, soil and decaying leaves (Naumov et al. 1998; Sniegowski et al. 2002; Zhang et al. 2010; Wang et al. 2012; Hyma and Fay 2013). Given their abundance in arboreal habitats, it seems unlikely that their exceptional fitness in grape juice is due to adaptation to grape juice fermentations and thereby provides evidence for exaptation (Larson et al. 2013). One way in which these species may have become adapted to high-sugar but low-nutrient environments is through associations with insect-honeydew, which is high in sugar (>10 g/l) but low in amino acids (Douglas 1993; Fischer and Shingleton 2001; Fischer et al. 2002). While a variety of insects and other animals exploit honeydew for its sugar resources (Beggs and Wardle 2006), and a recent investigation revealed that many taxonomically diverse fungi compete for honeydew (Dhami et al. 2013), no *Saccharomyces* species were found associated with aphid honeydew from Black Beach trees in New Zealand (Serjeant et al. 2008). It is also possible that *S. cerevisiae* is not adapted to a particular niche but is a generalist that happens to be particularly fit in high-sugar environments (Goddard and Greig 2015).

In addition to dominance relationships we studied, a variety of other factors may contribute to *S. cerevisiae*’s observed dominance of grape juice fermentations in vineyards. Many WGD species may not be present within vineyard environments. Of the non-*Saccharomyces* species used in this study, only the non-WGD species *Z. rouxii*, *T. delbrueckii*, *L. thermotolerans*, and *H. vineae* have been reportedly isolated from grapes or wine must (Kurtzman et al. 2011). Furthermore, sulfites are frequently added to wine must, and wine strains of *S. cerevisiae* are known to exhibit higher levels of sulfite resistance than non-wine strains (Pérez-Ortín et al. 2002; Yuasa et al. 2004). Interspecific differences in sulfite resistance (Engle and Fay 2012), copper resistance (Warringer et al. 2011) and other environmental conditions that we did not examine may thus contribute to *S. cerevisiae*’s observed dominance of wine fermentations. Although we find *S. paradoxus* to be competitive with *S. cerevisiae* in controlled laboratory settings, it has been reported to be the dominant yeast species in only a few fermentations (Redzepovic et al. 2002; Valero et al. 2007). Complicating such comparisons, however, is the possibility that *S. paradoxus* may not have always been distinguishable from *S. cerevisiae*, particularly in earlier studies of dominant yeast species from spontaneous wine fermentations.

## Acknowledgements

We thank members of the Fay lab for comments and suggestions. This work was supported by a National Institutes of Health training grant to KRW (HG000045) and research grant to JCF (GM080669).

## Supplemental Information

**Table S1. Yeast strains used in this study.**

**Table S2. PCR and pyrosequencing primers.**

**Table S3. PCR primers, barcodes and adaptors used for high-throughput sequencing.**

**Table S4. Significance of changes in the relative abundance of *S. cerevisiae* compared to other yeast species in high-sugar environments.**

**Table S5. Significance of intrinsic growth rate differences between *S. cerevisiae* and other yeast species in high-sugar environments.**

**Table S6. Significance of IC_50_ and intrinsic growth rate differences between *S. cerevisiae* and other yeast species in ethanol (%).**

**Table S7. Significance of intrinsic growth rate differences between each species’ own supernatant and supernatant from *S. cerevisiae* mono-culture and co-culture.**

**Table S8. Significance of low-pH on the intrinsic growth rate of *S. cerevisiae* and other species in HS.**

**Table S9. Significance of nutrient supplements on the intrinsic growth rate of S. *cerevisiae* and other species in Grape.**

**Figure S1.**
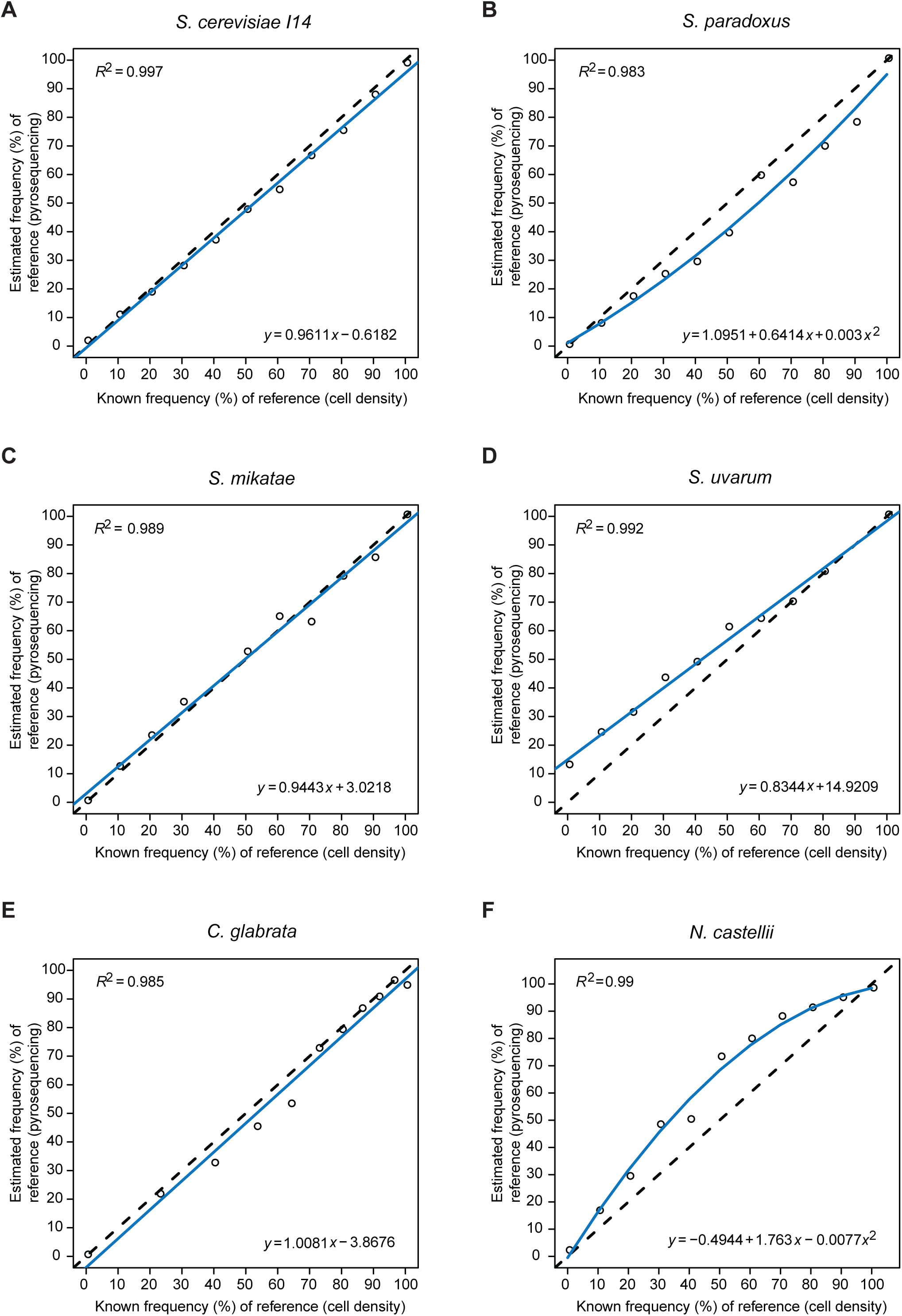
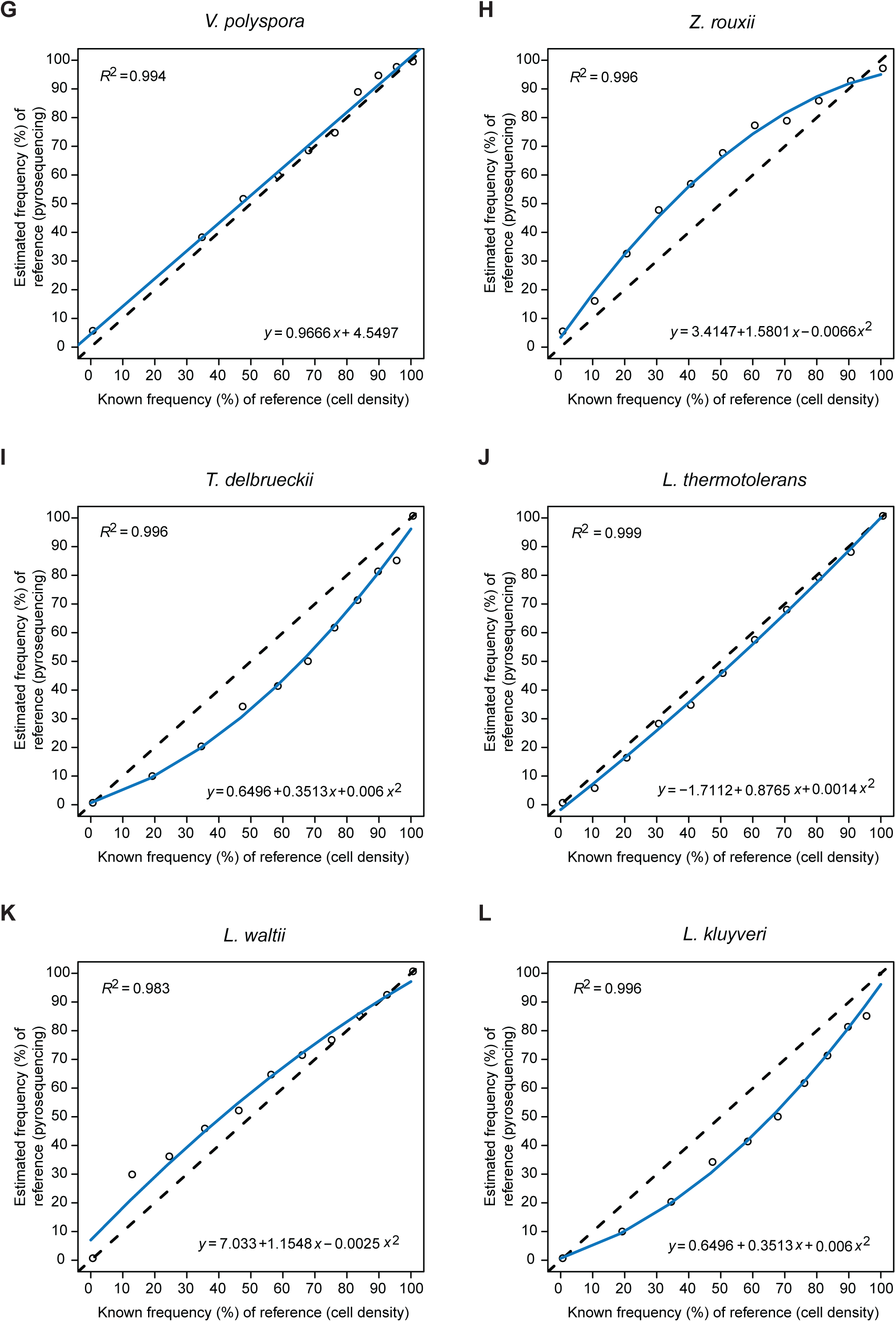
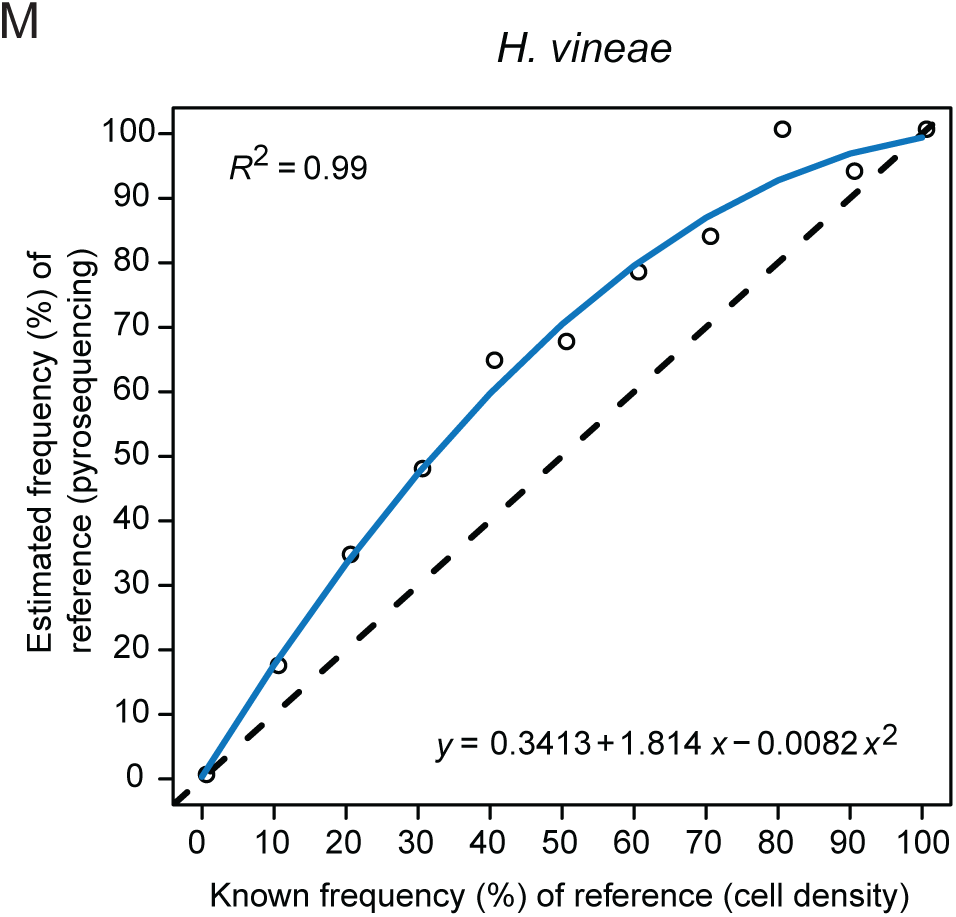
Pyrosequencing calibration. Relationship between the known frequency (%) of the reference species based on cell density and the estimated frequency (%) of the reference species based on pyrosequencing relative to *S. cerevisiae* (YPS163). Reference species include *S. cerevisiae* (I14) (A), *S. paradoxus* (B), *S. mikatae* (C), *S. uvarum* (D), *glabrata* (E), *N. castellii* (F), *V. polyspora* (G), *Z. rouxii* (H), *T. delbrueckii* (I), *L. thermotolerans* (J), *L. waltii* (K), *L. kluyveri* (L), and *H. vineae* (M). Calibration equations based on linear or polynomial regression analysis and *R*-squared values for each model are indicated.

**Figure S2.**
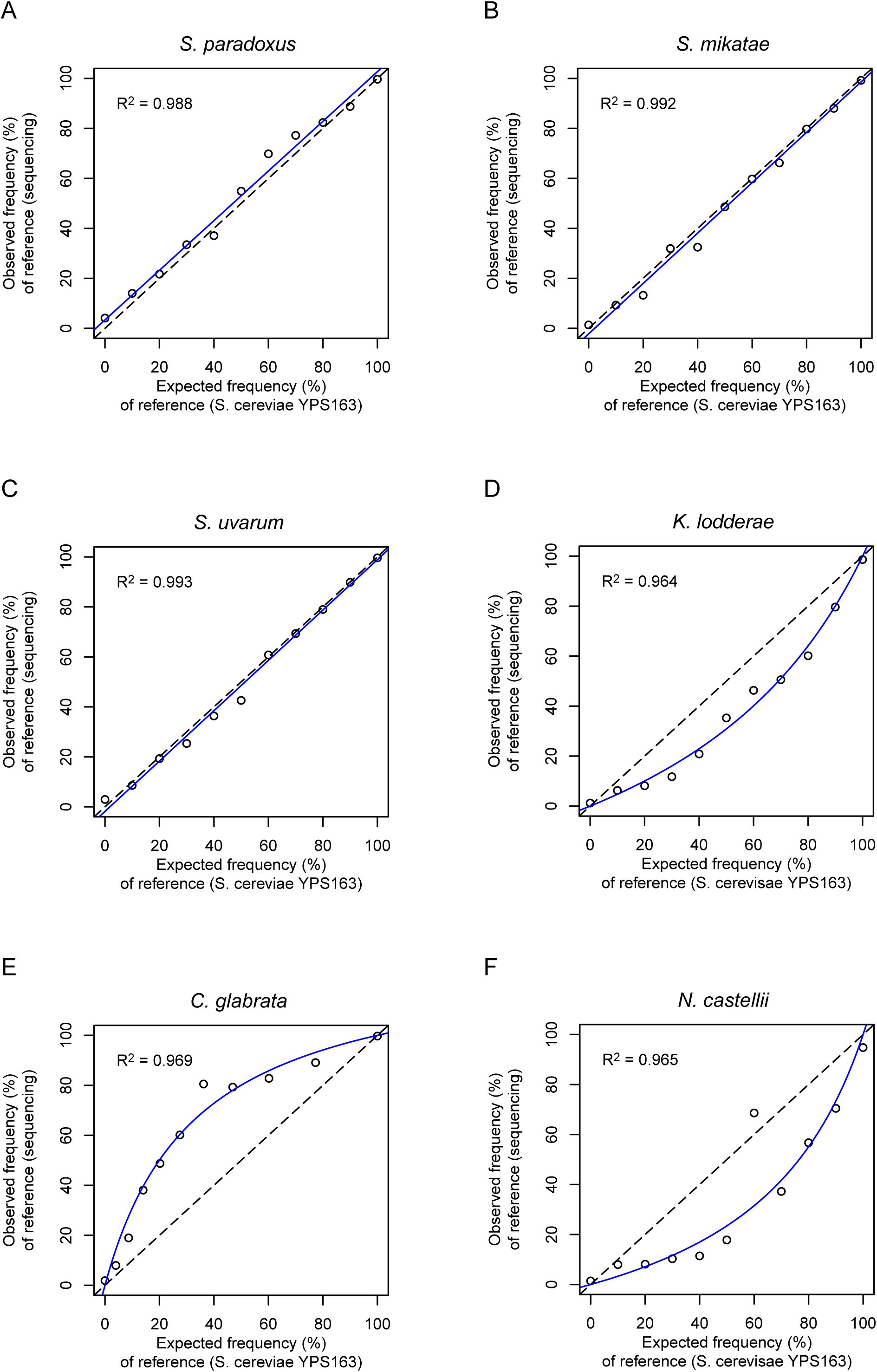
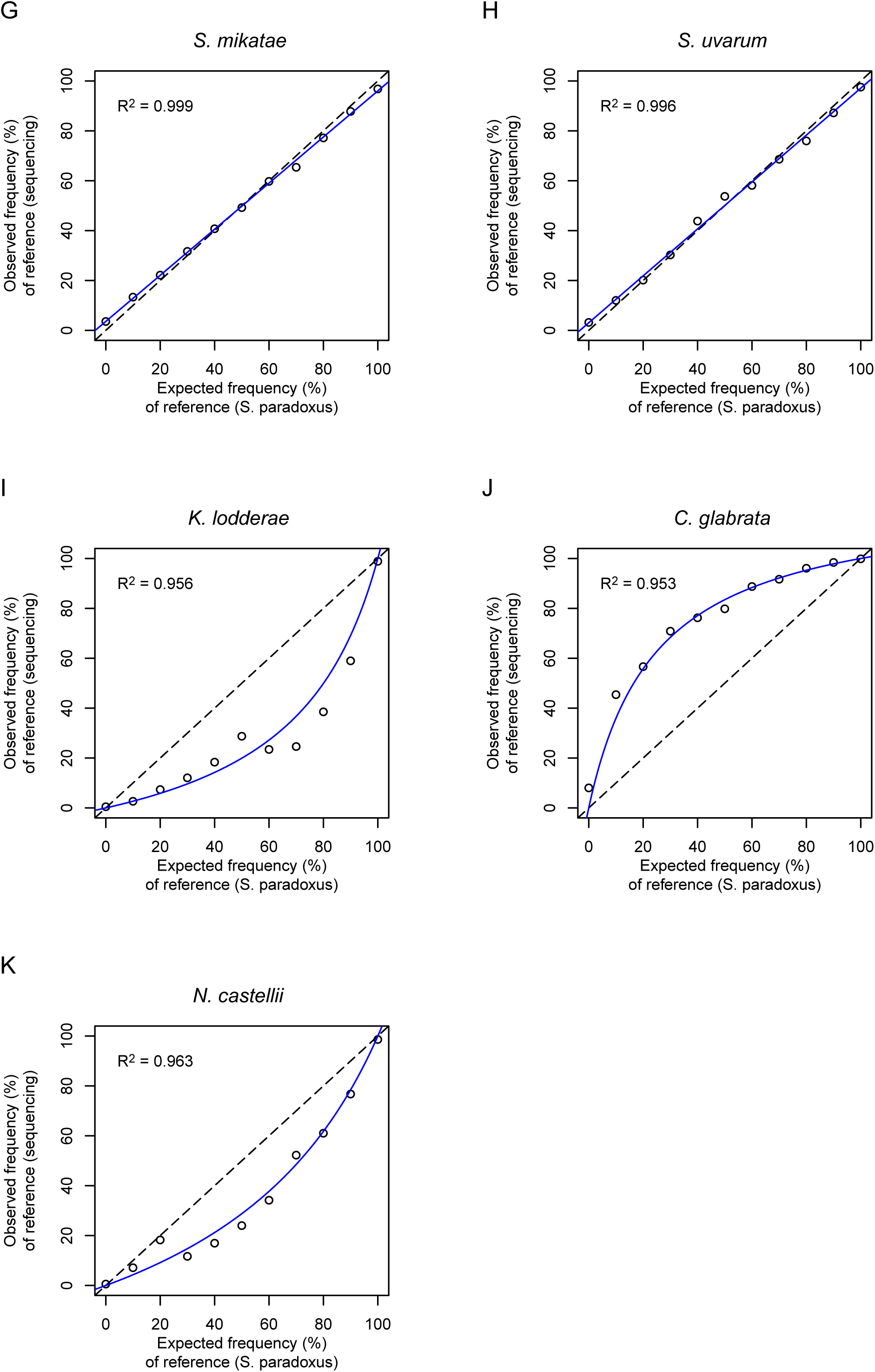
High-throughput sequencing calibration. Relationship between the known frequency (%) of the reference species based on cell density and the estimated frequency (%) of the reference species based on high-throughput sequencing. Panels show the relative abundance of the *S. cerevisiae* reference (YPS163) in comparison to *S. paradoxus* (A), *S. mikatae* (B) *S. uvarum* (C), *K. lodderae* (D), *C. glabrata* (E) and *N. castellii* (F); and the relative abundance of *S. paradoxus* (YPS152) in comparison to *S. mikatae* (G), *S. uvarum* (H), *K. lodderae* (I), *C. glabrata* (J) and *N. castellii* (K). *K. lodderae* and *N. castellii* calibrations with *S. cerevisiae* are based on ACT1; all others are based on ITS1. Calibration curves (blue lines) and *R*-squared values are shown in each panel.

**Figure S3.**
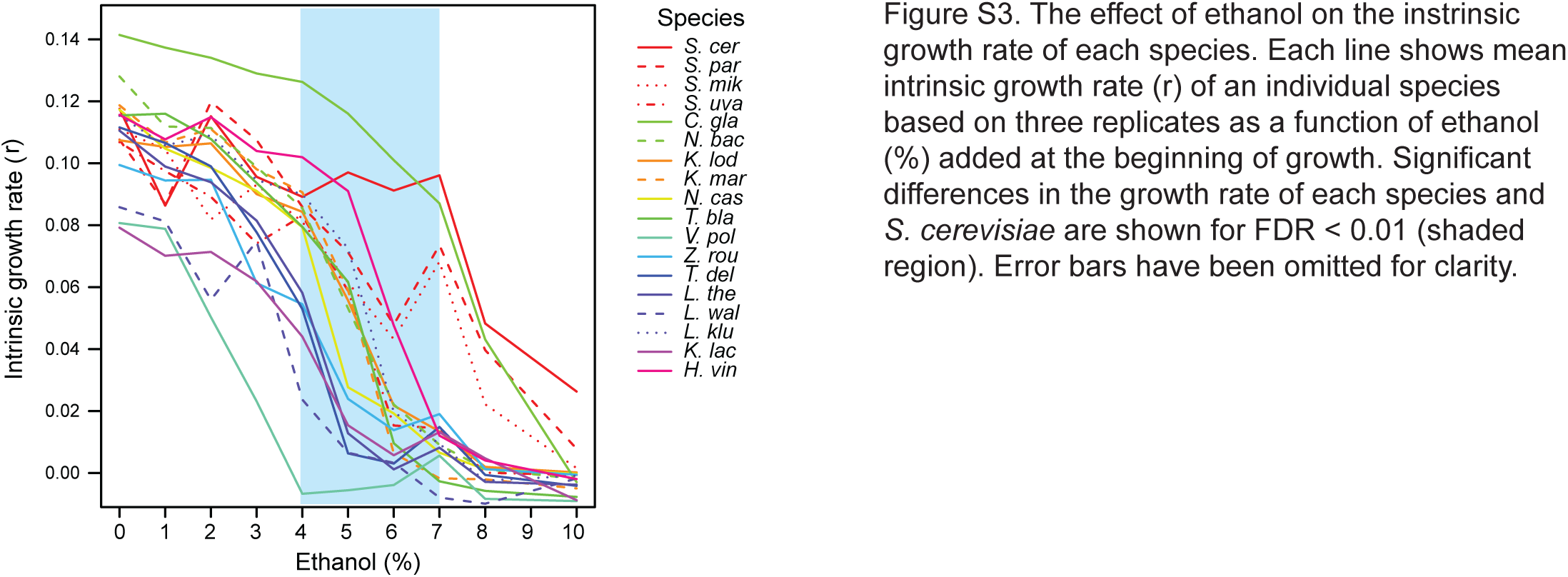
The effect of ethanol on the intrinsic growth rate of each species. Each line shows mean intrinsic growth rate (*r*) of an individual species based on three replicates as a function of ethanol (%) added at the beginning of growth. Significant differences in the growth rate of each species and *S. cerevisiae* are shown for FDR < 0.01 (shaded region). Error bars have been omitted for clarity.

**Figure S4.**
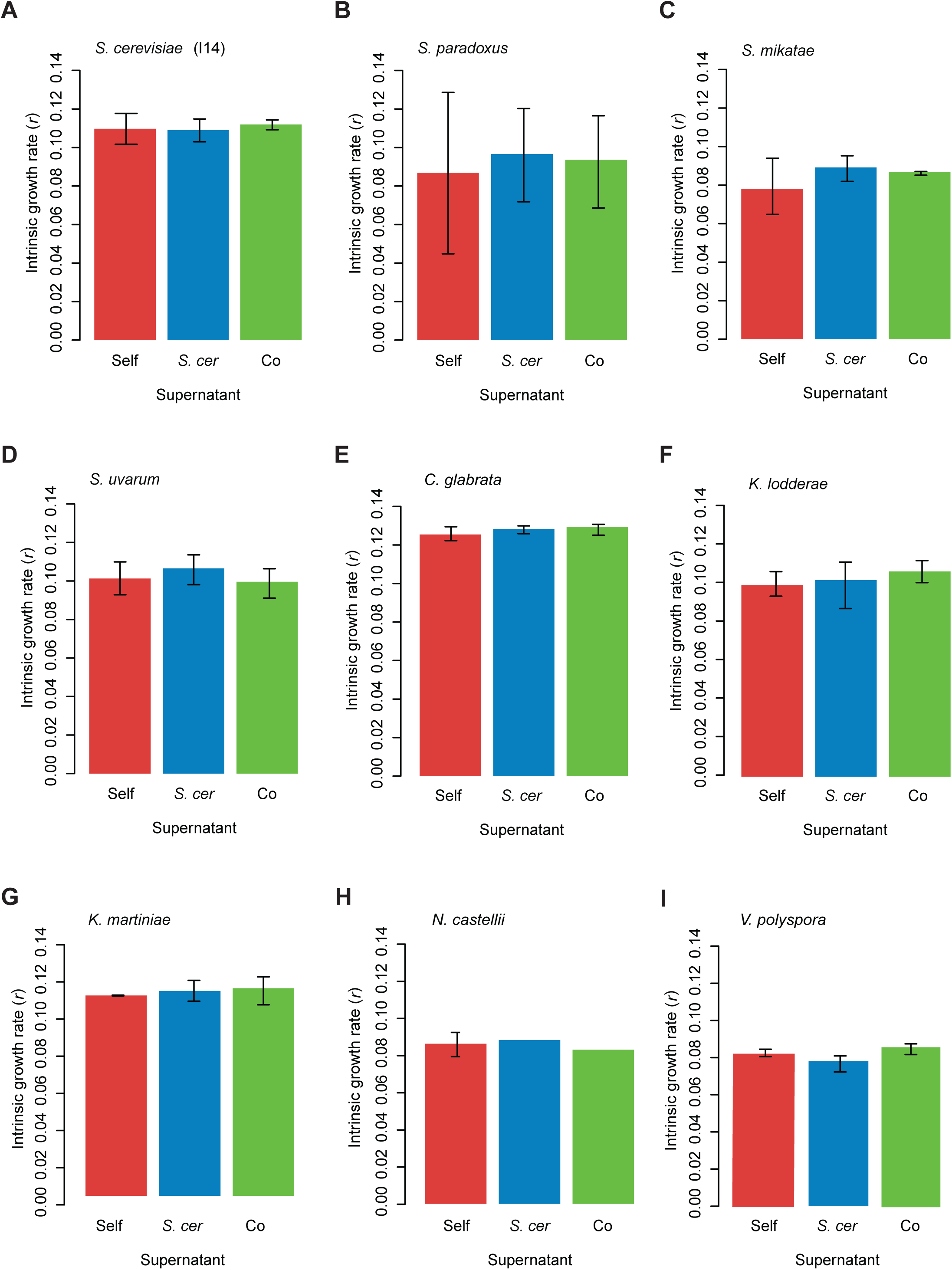
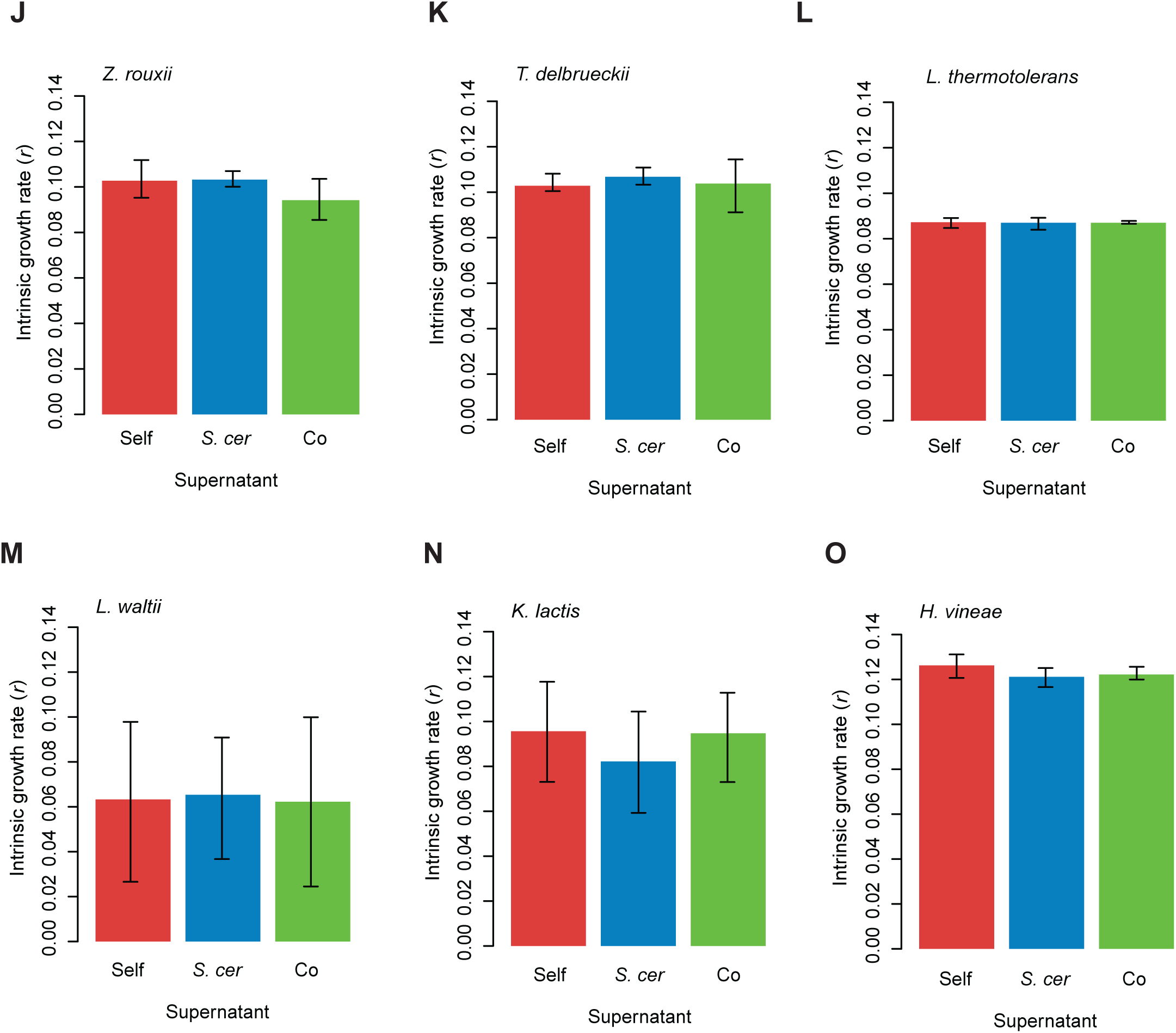
Intrinsic growth rate in YPD made with supernatant. The intrinsic growth rate of each species (A-O) in YPD made with the supernatant from each species’ own supernatant when grown in mono-culture (red), the supernatant from *S. cerevisiae* (YPS163) grown in mono-culture (blue), and the supernatant from co-culture with *S. cerevisiae* (green). Species are *S. cerevisiae* (I14) (A), S. *paradoxus* (B), *S. mikatae* (C), *S. uvarum* (D), *K. lodderae* (E), *K. martiniae* (F), *N. castellii* (G), *C. glabrata* (H), *V. polyspora* (I), *Z. rouxii* (J), *T. delbrueckii* (K), *L. thermotolerans* (L), *L. waltii* (M), *K. lactis* (N), and *H. vineae* (O). Bars and whiskers represent the mean and standard deviation of the growth rate.

**Figure S5.**
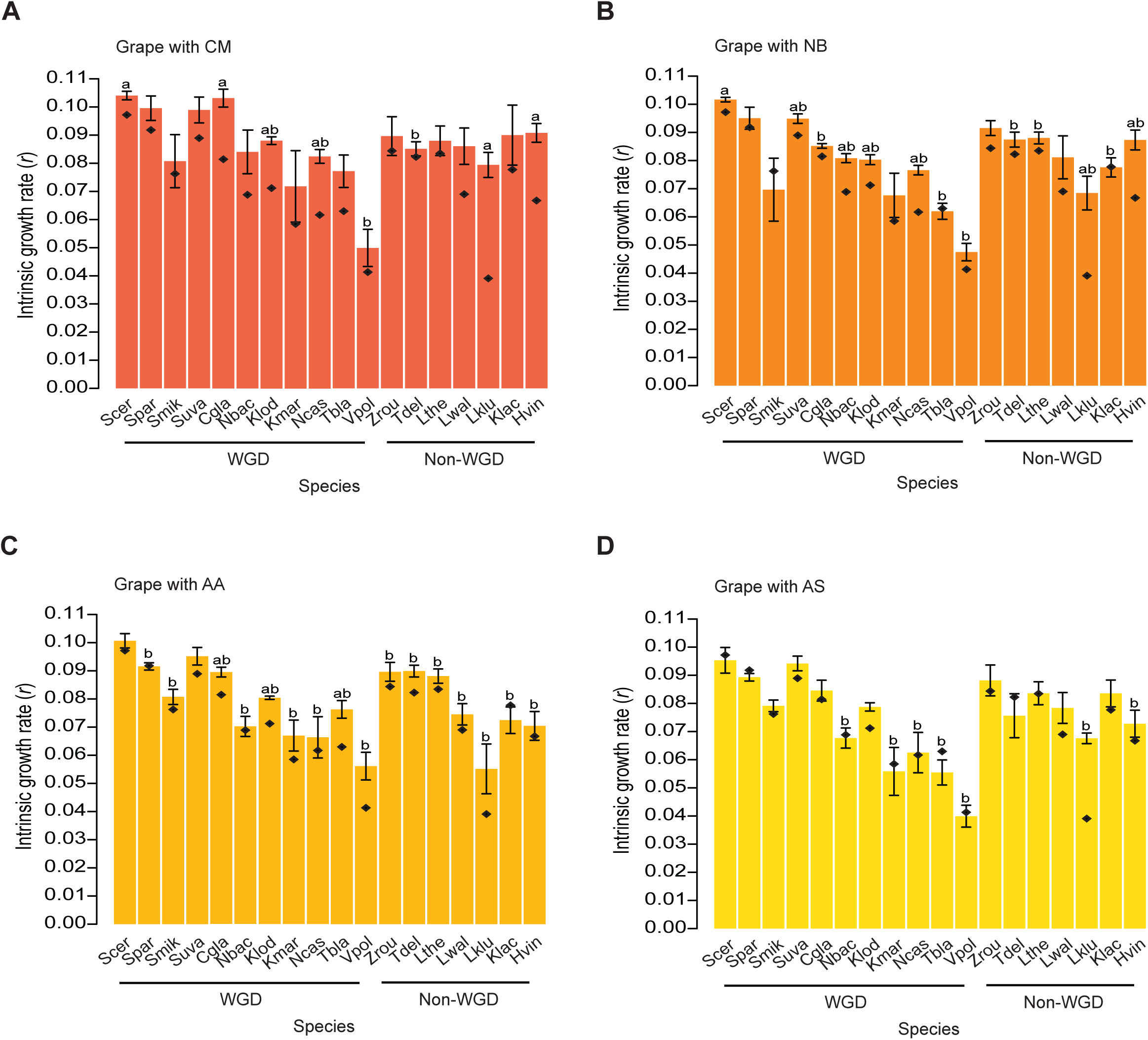
Intrinsic growth rate in Grape supplemented with various nutrients. The mean intrinsic growth rate of each species in Grape supplemented with CM (A), NB (B), AA (C) and AS (D). Names of WGD (blue) and non-WGD (red) species are colored. Bars and whiskers represent the mean and standard deviation of the growth rate. Significant differences in the growth rate of each species with or without the added nutrient (a) and differences between *S. cerevisiae* and each species with the added nutrient (b) are labeled above each bar for FDR < 0.01.

